# Influence of Non-Specific Surface Adhesion on the Shape and Microrheology of Red Blood Cells

**DOI:** 10.64898/2026.06.23.734082

**Authors:** Agathe Nidriche, Delphine Débarre, Claude Verdier

**Affiliations:** Université Grenoble Alpes, CNRS, LiPhy, 38400 Grenoble, France

## Abstract

Poly-L-Lysine (PLL) mediates the non-specific adhesion of cells and is commonly used in Atomic Force Microscopy (AFM) measurements, to ensure that cells remain attached to the substrate. However, it is acknowledged that adhesion affects the measured mechanical properties, in particular in the case Red Blood Cells (RBCs). This results in a wide range of Young’s modulus *E* reported in the literature. The present study aims at providing a systematic approach to the impact of non-specific adhesion on the rheology of RBCs. It provides a correlation between the topography profile of adherent RBCs and their rheology, from weak (*c*_PLL_ = 10^−3^ mg/mL) to strong-adhesion (*c*_PLL_ = 10^0^ mg/mL) regimes. Using RICM and AFM, we find that there is a continuum of RBC shapes promoted by adhesion, from concave to dome-shaped, as predicted by the theory of vesicle adhesion. Their elastic properties discriminate them into two populations depending on adhesion strength, where stiffer RBCs (*E* ≳ 100 Pa) correlate with dome-shaped cells. These findings are supported by rheology measurements of the dynamic complex shear modulus *G*^∗^(*f*): while the storage modulus increases with cell-substrate adhesion, reflective of an increased membrane shear modulus, the loss modulus remains unchanged. Finally, further analysis inspired by membrane theory shows that different deformation modes may be triggered during indentation of either weakly or strongly adhering RBCs, illustrating the limits of the Hertz model.

## Introduction

Red blood cells (RBCs) constitute about half of the blood content and their extreme deformability allows them to pass through submicrometer channels [1], which is responsible for the particular viscoelastic properties of blood [2]. Their cholesterol-rich membrane has an extremely low bending modulus, which accounts for their native biconcave-shape. A quasi-hexagonal prestressed 2D spectrin lattice is anchored in the membrane through different protein complexes. RBCs are responsible for O_2_ exchanges in blood, hence their cytosol is mainly constituted of hemoglobin which is responsible for its viscosity. RBCs are deprived of organelles, and for that reason were long considered as passive cells due to their lack of active motility and mitochondrial metabolism. However, recent studies proved functional activity in the RBC. A first example is the mechanosensitive channel Piezo 1 regulating calcium entry and cell volume [3], and another one is the necessity to account for ATP-driven phosphorylation of spectrin and membrane anchorage proteins such as Band 4.1, as well as ion pumps activity, to rationalize the non-equilibrium membrane fluctuations at low frequencies *f <* 10 Hz [4, 5]. Yet, RBCs remain a typical minimalistic model for a non-motile cell.

For that reason, RBCs have been studied for decades, and seminal works consisted of describing their behaviour through the lens of elastic membrane mechanics [6, 7], accounting for the main elastic deformation modes provided by the membrane and cytoskeleton [8]. In the 1980’s, Evans, Skalak *et al.* provided values for their bending (*κ* ≈ 1.8 10^−15^N/m [9]), shear (*µ* ≈ 7 10^−6^ N*/*m [10]) and area dilation (*K* ≈ 4.5 10^−1^ N*/*m [11]) elastic moduli using micropipette experiments [12]. These authors further described the viscoelastic properties of RBCs and provided estimations for the membrane viscosity (≈ 10^−6^ Ns*/*m [10]). Early works also estimated the Young’s modulus *E* from the relaxation times of RBCs undergoing hemolysis [13], micropipette expel-lation [14] and uniaxial stretch [15] to be in the range of 1 − 10 kPa. *E* is shown to increase with storage time [16], and is expected to be dependent on cell shape [17] or pathologies [18]. The emergence of other nanoscale techniques such as Atomic Force Microscopy (AFM), optical tweezers [19], microfluidics [20] or even neutron spin echo [21] coupled with simulations [5] further expanded the knowledge of single RBCs mechanics, while opening more questions about their multiscale dynamics and behaviour in flow and constrictions.

Studies of the viscoelasticity of the RBCs estimate their complex shear modulus *G*^∗^(*f*) over a range of frequencies *f*. Passive rheology of the membrane’s fluctuations has been carried-out with Dynamic Scattering Microscopy (DSM) [22] and diffraction phase microscopy [23]. Otherwise, active rheology measurements were performed using force exerting methods, as for instance applying a torque to a bead attached to the membrane with Magnetic Twisting Cytometry (MTC) [24], stretching the cell in between beads using optical tweezers [25], or using an acoustic force microrheology method (AFMR) [26]. All these methods provide a cut-off frequency *f_t_*≈ 10 Hz defined by *G*′(*f_t_*) = *G*′′(*f_t_*) where *G*′ and *G*′′ respectively denote the storage and shear moduli obtained from *G*^∗^. *G*_0_, the storage modulus at low frequencies, is found in a large range of values *G*_0_ ∈ [10^−2^, 10^2^] kPa. AFM is also a key technique for the description of intracellular viscoelasticity [27]. Yet, it has been used mainly to describe the elastic properties of RBCs, either using the formalism of membrane mechanics [28] or considering that the RBC is isotropically elastic to obtain an effective Young’s Modulus *E*_eff_. However, measurements for *E*_eff_ span a few orders of magnitude from 100 Pa [29, 30] to 100 kPa [31, 32], and do not converge to a consensus value. An impediment of AFM measurements is the necessity that cells adhere to the substrate, to avoid additional structural deformation. Among common immobilizing methods are microwells, patterning, or tuning the surface properties of the substrate on which cells are deposited. An ubiquitously-used mediator of cell-substrate adhesion is Poly-L-Lysine (PLL), a polyelectrolyte which is positively charged at pH 7 and which promotes non-specific adhesion through electrostatic interaction with the negatively-charged RBC glycocalyx. Hategan *et al*. [33] have shown that PLL concentration and molecular weight control the emergence of adhesive patterns and lipid domains formation in RBC, where Band 3 protein and lipids are in direct interaction with PLL, and provide an insightful phase diagram of this topographical patterns at the surface of the substrate. They show that a PLL concentration of *c*_PLL_ = 10 mg*/*mL ensures a strong RBC adhesion within the fraction of a second, inducing a spherical-shape geometry of the RBC with robust adhesion area *A* ≈ 60 *µ*m^2^ and contact angle *θ* ≈ 60^◦^.

However, experiments requiring cell adhesion are usually performed in the lower adhesion regime for a wide range of concentrations from *c*_PLL_ ≈ 10^−4^ to *c*_PLL_ ≈ 1 mg*/*mL, where membrane bending is non-negligible. It is unclear how and to what extent substrate adhesion impacts AFM measurements. Controlling the shape of the RBC both at the substrate and in contact to the AFM tip is paramount for force-indentation curves analysis. Yet, most AFM measurements are performed without monitoring precisely the PLL concentration.

This study aims at tackling the lack of experimental information regarding the shape transition of a single RBC upon variation of non-specific substrate adhesion strength, and quantifying how it relates to its measured dynamical mechanical properties. For that purpose, we probe a large range of PLL concentrations from 10^−3^ to 1 mg*/*mL at fixed molecular weight, locally covering 1% to 90% of the substrate’s surface. The change in membrane curvature triggered by substrate adhesion is probed using RICM and AFM, and correlated to the effective stiffness of RBCs. Modifications in shape arising from substrate-adhesion also highlight the impact of membrane deformation on force-indentation curves, and draw a very narrow range of applied forces where standard linear models can be applied. In that regime, a detailed study of RBC rheology illustrates how the complex shear modulus *G*^∗^(*f*) varies with adhesion.

## Materials and Methods

### Red blood cells preparation

Human blood was collected at the Etablissement Français du Sang (EFS Rhône-Alpes Auvergne, France) and stored at 4°C with citrate anticoagulant in aliquots of volume V=2.7 mL. These samples were used within 3 days after collection. A 0.5 mL volume of blood was resuspended in an isotonic solution of 1.5 mL 1X PBS (PBS tablets 10 mM Phosphate, 2.68 mM KCl, 140 mM NaCl, pH = 7.45, Gibco) and then centrifuged at 6000 rpm (Sprout mini, Thermofisher) and resuspended 3 times to isolate erythrocytes.

Poly-L-Lysine of molecular weight 70 − 150 kDa (P1274-500MG lyophilised powder, Sigma-Aldrich) was stored at −20^◦^C and diluted before use at concentrations of 0.001 mg*/*mL,0.01 mg*/*mL, 0.1 mg*/*mL and 1 mg*/*mL. A polystyrene plastic plate (TPP AG, Trasadingen, Switzerland) with a glued borosilicate glass coverslip (VWR International, Radnor PA, USA, 631 − 0175) was thoroughly rinsed with ethanol and distilled water before use, and air-plasma treated for 10 s to induce better PLL adsorption. Following the manufacturer protocol, 500 *µ*L of Poly-L-Lysine was deposited on the plate for 10 minutes, before being rinsed with sterile ultrapure water (resistivity 18.2 MΩ) and let to dry for a minimum of 2 hours under the sterile bench to allow the formation of a secondary structure [34]. 2 mL of a suspension of freshly rinsed RBCs at a concentration of approximately 0.1 g*/*L was deposited on the coated Petri dish for 5 minutes. In order to keep only adherent RBCs and avoid diffusion-induced noise during measurement, the medium was gently removed and replaced with 2 mL of PBS. After temperature equilibration, measurements were performed for a maximum of 40 min before spontaneous adhesion-induced RBC lysis (43 min characteristic time for *c*_PLL_ = 10 mg*/*mL, Hategan et al [34]).

### PLL coating characterization

In order to estimate the amount of adsorbed PLL as a function of the solution concentration, the thickness of the dried adsorbed layer on silicon wafers was measured with ellipsometry, with *c*_PLL_ bulk concentrations ranging from 10^−2^ mg*/*mL to 10 mg*/*mL. To this aim, silicon wafers were air plasma-cleaned for 3 minutes before deposition of 1 mL of PLL solution of concentration *c*_PLL_ on the wafer placed in a 30 mm diameter glass dish. After 10 min incubation, the PLL solution was removed and the wafer was rinsed 3 times with sterile ultrapure water, and left to dry under sterile conditions for 6 hours. Ellipsometry measurements were performed on a rotating quarterwave plate home-built instrument [35]. The linearly polarized laser beam with incident wavelength *λ* = 633 nm worked at a 70^◦^ grazing angle. Data analysis assumed a double-layer model including a first silicon layer covered by a second SiO_2_+PLL layer of refractive index 1.46, which height is to be determined. Beforehand, the height of the oxidized layer 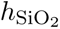 was measured on the clean silicon wafer. After functionalization, the PLL layer thickness was obtained by subtracting the oxidized silicon layer thickness to the total measured thickness. Each wafer was measured at 10 different spots.

The mean height of the PLL layer was modeled using a Langmuir’s adsorption isotherm, see Figure S2, accounting for the deposition of a single layer of non interacting particles until a plateau of adsorption is reached. This yields for the average height *< h*_PLL_ *>*

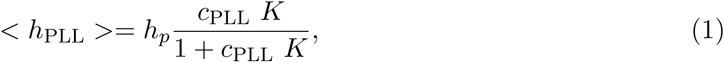

where *h_p_*= 0.8 nm is the plateau PLL height at full coverage and *K* = 9.2 mL*/*mg is the adsorption coefficient. The coverage is defined by 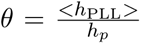, and estimated by the model is presented in Table 1.

**Table 1:** PLL layer average height *h*_PLL_ and corresponding coverage *θ* as a function of PLL solution.

|  |  |  |  |  |
| --- | --- | --- | --- | --- |
| $c_{\text{PLL}}$ [mg/mL] | $10^{-3}$ | $10^{-2}$ | $10^{-1}$ | 1 |
| $h_{\text{PLL}}$ [nm] | 0.01 | 0.07 | 0.40 | 0.76 |
| $\theta$ [%] | 1 | 10 | 50 | 90 |

In addition to ellipsometry measurements, the uniformity of the PLL layer was qualitatively assessed using an inverted microscope (Zeiss observer Model D1 Jena, Germany) in epifluorescence mode: to this aim, PLL solutions at different concentrations were fluorescently labelled using sulforhodamine B LC 6200 (Lambdachrome laser dye, Lambda Physik), introduced in stoe-chiometric molar concentration right before a 10 minutes incubation on the bottom plate. Images with a 0.271×0.169 *µ*m field of view were scanned in steps of ≈ 0.5 *µ*m, to assess the uniformity of the PLL layer across the whole bottom-plate. Following a calibration of PLL concentration with fluorescence intensity (Figure S1 a), it is observed that the mean fluorescence intensity varies with a standard deviation of 34 ± 12% over a single plate (Figure S1 b). It illustrates the necessity to probe RBCs over the whole plate within a single experiment, in order to obtain statistically representative results.

### Reflection Interference Contrast Microscopy

Reflection Interference Contrast Microscopy (RICM) was used to decipher the topography of the cell adhering region for the different PLL concentration conditions used for AFM measurements. It also assessed the uniformity of the adhesion pattern, as well as the presence of thermal fluctuations of the RBC membrane. The set-up was described previously in Ref. [36].

Using previously-described PLL surface coatings, RBCs were seeded directly under the microscope with a concentration found to yield a few dozen separated cells per frame. In the case of *c*_PLL_ = 0 mg*/*mL, the plate was also treated with air-plasma. Interference patterns were probed with 3 different wavelengths (*λ* = 453, 532 and 632 nm), and compared with bright field microscopy. Images with a 0.108 *µ*m*/*px sampling and a lateral resolution approximately 250 nm were captured at *T* = 37 ^◦^C with the focus either on the substrate’s surface to assess the adhesion pattern and using optical sectioning [37], or at the top of the cell to assess the upper shape of the RBC, see Figure 1. For membrane fluctuation analysis at the top part of the cell, sequences of 100 images with a 100 ms frame interval were acquired, and the coefficient of variation of membrane fluctuations *CV* = 100(⟨*δI/*⟨*I*⟩⟩ − ⟨*δI/*⟨*I*⟩⟩_background_) was measured within each cell (*N* = 519 cells) with a custom-made routine using Fiji [38]. Since adhesion and spreading on the coated coverslip occured in less than a second for *c*_PLL_ *>* 10^−1^ mg*/*mL, measurements of the adhesion area were performed using a single image per cell, which is considered representative of its adhesion state during mechanical measurements.

**Figure 1:**
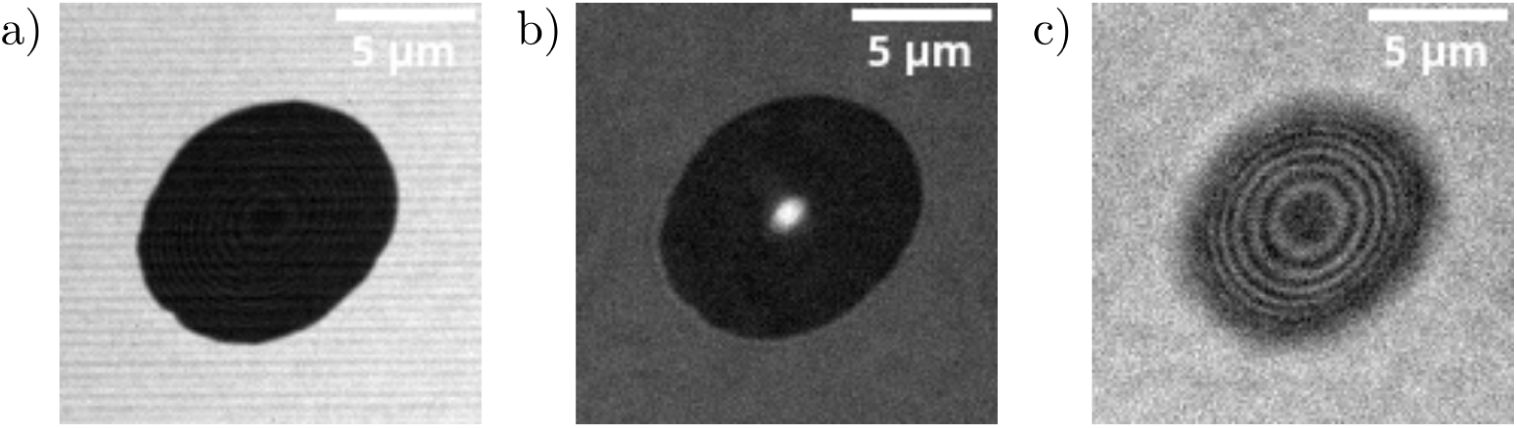
Different acquisition used for the study of RBC adhesion with RICM, *c*_PLL_ = 1 mg*/*mL, *λ* = 532 nm. a) Focus on the substrate, using optical sectioning with structured illumination. Typical image used for cell-substrate contact area segmentation. b) Focus on the substrate without optical sectioning. The central bright spot is due to the reflection from the domed-shape upper surface of the RBC. c) Focus on the top of the RBC, no optical sectioning. Typical image used to study the fluctuations of the upper membrane.

Segmentation of such images was performed using Fiji. The whole image stack was corrected for the background with a pseudo flat-field correction (dividing by the same Gaussian-blurred image with a s.t.d. of 1*µ*m), normalized, then thresholded using the MinErr method over the whole stack. Cell counting was performed with the “analyze particles” plugin with a circularity cutoff of 0.6 and a contact area range *A*^∗^ ∈ [3 − 120]*µ*m^2^.

### AFM topography

Simultaneous high-speed scanning of height and force-indentation curves was performed on live cells maintained at *T* = 37 ^◦^C using an atomic force microscope (NanoWizard 4XP, Bruker, Germany) working in QI™Advanced Imaging mode, with optimized cantilever and sample motions to acquire high-quality images. The AFM is coupled with an inverted microscope (Zeiss observer Model D1 Jena, Germany, ×10 and ×40 objectives). Precalibrated PFQNM-LC-V2 cantilevers (Bruker, Germany) of cylinder-shaped geometry with stiffness *k* ≈ 0.15 N*/*m were used. Scans were performed with a 36 px × 36 px resolution in a square of dimensions 14 *µ*m × 14 *µ*m, and acquired at speed 70 *µ*m*/*s, with a force setpoint *F* ≈ 50 pN. To fit force-indentation curves, we used the model of a flat cylinder of radius 70 nm, corrected for the effect of the substrate [39]. The sample height was estimated for each pixel using the contact point of the approach force curve, obtained with the cylinder model for *F* ≤ 30 pN. The corresponding vertical tip displacement provides the indentation of the tip in the sample. A custom Python script was used to correct measurement artefacts, subtract the background and estimate the radially-averaged height and indentation profile of the RBCs for all PLL concentrations, assuming a radially symmetric height profile.

### AFM microrheology measurements

Precise mechanical measurements (1 force-indentation curve per RBC, measured at the center) were performed using the same AFM setup, but different cantilevers and indentation parameters. Spherical beads of 3 *µ*m diameter (Fluoresbrite®YG Carboxylate Microspheres Polysciences) were glued with epoxy resin onto tipless triangular shape cantilevers with stiffness *k* = 0.02 N*/*m (QUEST T 20, Nunano, Bristol, UK). The position and presence of the bead were assessed using fluorescence imaging before experiments, which also permitted precise positioning of the bead at the center of the RBC. The carboxylate beads were treated with 0.5% BSA in PBS to reduce tip/cell adhesion.

For each blood donor (11 in total) and each PLL concentration, 20 RBCs per PLL concentration were measured a single time at the center. We verified that repeated measurements over a single cell are reproducible (s.e.m. ≤ 1 Pa, measured for *N* ≈ 15 curves/cell for each studied concentration), and that the cell is not stiffening over time.

The sample was indented with a force setpoint *F* = 30 pN.The resulting distribution of indentations *δ*_0_ measured across all cells spans from the first quartile *Q*1 = 250 nm to the third quartile *Q*3 = 900 nm, depending on the cell’s elastic properties (see Figure S4 (b)). Following the approach of the tip with velocity *v* = 1*µ*m*/*s, sinusoidal oscillations (5 periods) were imposed with fixed peak-to-peak amplitude *a* = 100 nm (for soft RBCs, *c*_PLL_ *<* 1 mg*/*mL) or *a* = 20 nm (for more rigid RBCs, *c*_PLL_ = 1 mg*/*mL). Frequencies in the range *f* ∈ [0.5, 100] Hz were successively probed using 3 values/decades. Measurements were performed one after the other for a single initial indentation *δ*_0_, and separated by 20 ms pauses at constant height.

For the analysis of the experimental data, the mechanical parameters used to describe the RBC mechanics were the Young’s modulus *E*, characterizing the elasticity of the RBC in the hypothesis of an isotropic elastic material using the approach curve only; and the complex shear modulus *G*^∗^, whose real and imaginary parts account for stored and dissipated energy. Both are determined within the framework of the Hertz model for the indentation of a semi-infinite flat cell by an infinitely rigid sphere [40, 41]. Knowing the initial indentation *δ*_0_ corresponding to the initial force setpoint, the complex shear modulus which is a combination of the storage modulus *G*′(*ω*) and the loss modulus *G*′′(*ω*) writes

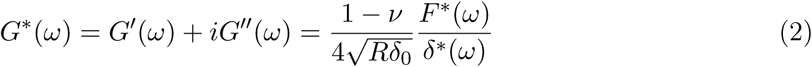

Where *F* ^∗^(*ω*) is the indentation force measured by the cantilever deflection, *δ*^∗^(*ω*) the height imposed by the piezo-actuator, *R* the radius of the spherical indenter, and it is assumed that the Poisson’s ratio of the incompressible RBC is *ν* = 0.5.

Motivated by the large indentation depth *δ*_0_ relative to the thickness of the RBC *h* ≈ 2 *µ*m (Figure S3,S4), the Young’s modulus *E* was corrected for the underlying hard substrate using Dimitriadis *et al.* correction factor 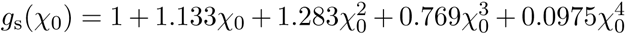 [42] for a spherical indenter of radius *R* = 1.5 *µ*m, yielding the corrected modulus *E_c_*

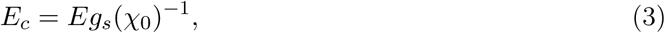

where 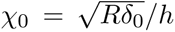. The detailed approach for the determination of *χ*_0_ is provided in the Supplemental Materials (Figures S3 and S4).

This correction factor *g*_s_(*χ*_0_) is generalized to yield the corrected complex shear modulus 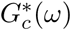 following the method exposed in Ref. [43, 44], assuming a negligible oscillating indentation *δ* ≪ *δ*_0_, and performing a Taylor expansion around initial indentation *δ*_0_:

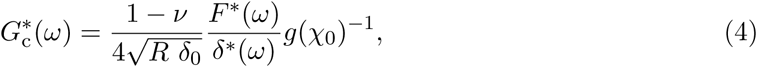

where *g*(*χ*_0_) is a generalized correction factor:

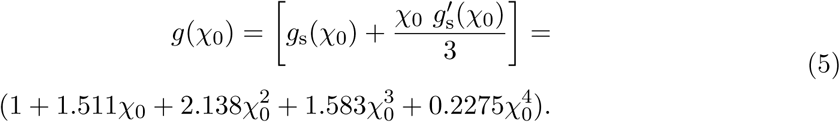

Further correction is performed for the hydrodynamic drag exerted by the PBS solvent on the tip in close contact with the sample. More details are provided in the Supplemental Materials. It is performed in a custom script using the procedure described in Alcaraz et al [45] with tip-substrate height spanning from *h* = 500 ± 50 nm to *h* = 5, 000 ± 50 nm and sinusoidal oscillations of peak-to-peak amplitude *d* = 300 nm measured for *f* ∈ [0.5, 200] Hz. The drag coefficient at contact is *b*(0) = 3.88 ± 0.03 *µ*Ns*/*m (see Supplemental Materials, Table S2 and Figure S6). The hydrodynamic drag exerted by the PBS solvent only affects the loss modulus, such that the complex shear modulus expressed in equation (4) and corrected for the hydrodynamic drag 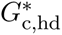 writes

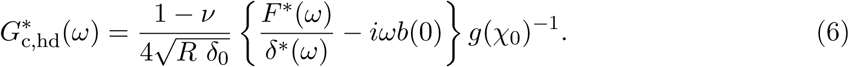

### Statistical analysis

Blood samples from different donors have been used. The total number of RBCs and donors measured for each experiment is detailed in Table S3. Statistical tests on data (fit parameters presented in Figure 6, indentation and substrate correction *χ* presented in Figure S4) were performed on Python, using a non-parametric Mann-Whitney-Wilcoxon test on ranks and corrected for multiple tests using a two-sided Holm-Bonferroni correction, using the “statannotations” library. Significance: ****: *p <* 10^−4^, ***: 10^−4^ *< p <* 10^−3^, **: 10^−3^ *< p <* 10^−2^, *: 10^−2^ *< p <* 5.10^−2^, ns: *p >* 0.05. Boxplots and violinplots feature the median, Q1 and Q3 values along with the distribution of datapoints, and are plotted using Python’s “seaborn” library. Rheology measurements were plotted using the median of data points for each concentration and the interquartile range [Q1-Q3] for the error bars. On Figure S9 c) and d), the analysis of force curves was plotted as the mean ±95% confidence interval (Student’s t-distribution with N-1 degrees of freedom), in order to display the variability of the individual curves. K-mean clustering and the quantification of intra-cluster cohesion and inter-cluster separation (silhouette score, *S* ∈ [−1, 1] from worse to best classification) are performed using scikit learn library on Python.

## Results and Discussion

### Non-specific cell-substrate adhesion triggers a transition in RBC geometry

The impact of cell-substrate adhesion on the overall shape of RBCs is studied both at cell-substrate interface using RICM, and at the upper membrane using RICM and AFM. Non-specific adhesion is modulated by varying *c*_PLL_, the concentration of Poly-L-Lysine ensuring a plate coverage from 1% to 90%.

RICM provides geometric and dynamic information on cell adhesion [46]. At *c*_PLL_ = 0 mg*/*mL for an air-plasma treated surface, a large proportion of RBCs display deformed circular rings with a large dark region at the cell-substrate interface. It is a signature that the RBC already shows non-specific adhesion but still remains biconcave and is only attached by its rim, with a cell-substrate distance in the range [0.3, 1.2] *µ*m. A typical observation is provided in Figure 2 (a). Strikingly, the adhesion of the RBC is complete already at 1% plate coverage: RBCs display a flat dark surface of adhesion area *A*^∗^ and a negligible proportion of incomplete adhesion, see Figure 2 (a). Hence, the biconcave shape is not conserved with substrate adhesion: this justifies the further assumption that cells are fully adherent and that structural deformation is negligible for AFM indentation measurements. It shows that the work of adhesion exceeds the theoretical limit *W_a_* = 2*κ/R*^2^ [47], with *κ* the bending rigidity of the RBC.

**Figure 2:**
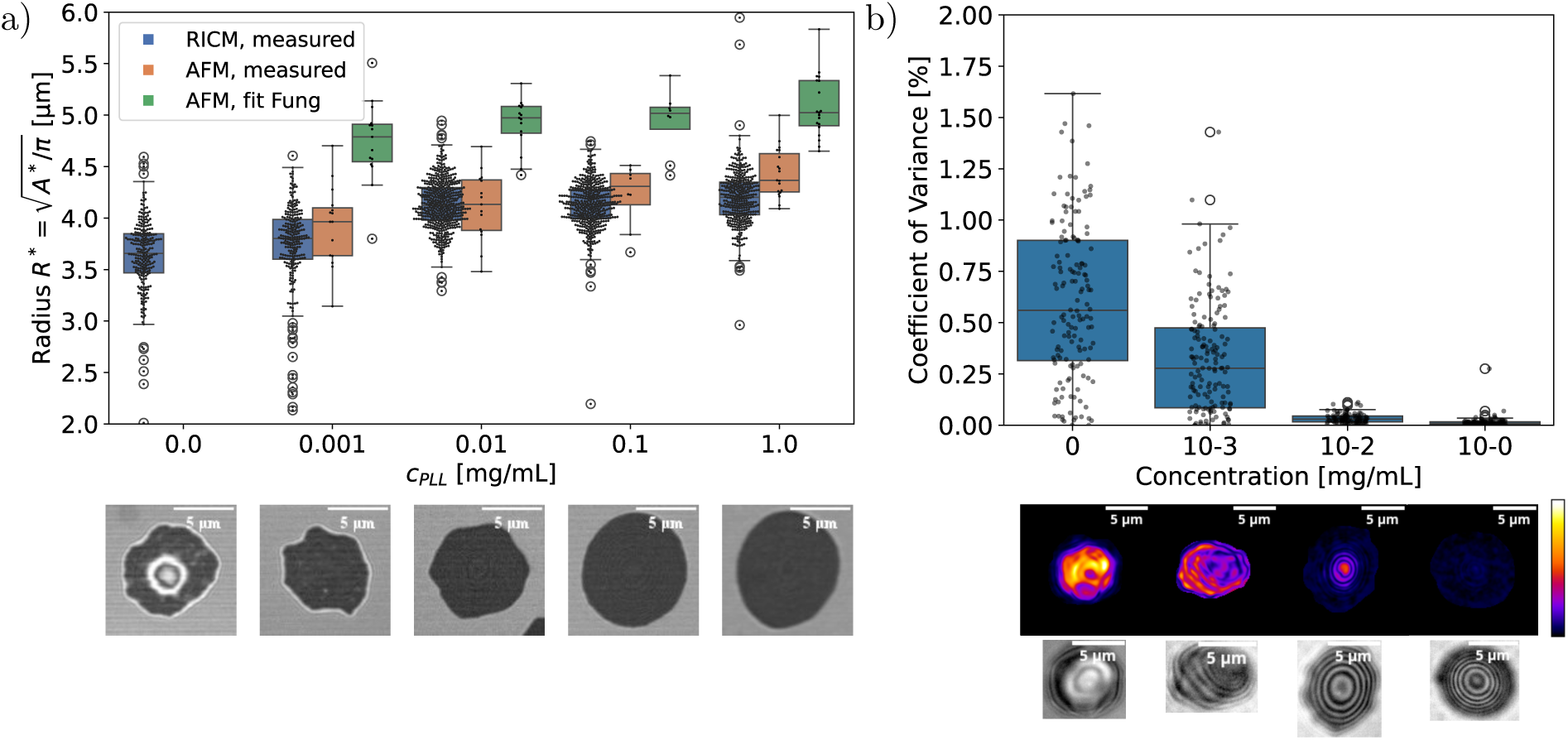
Bottom surface description of the RBC: a) Radius 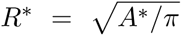 of the RBC in direct contact with the substrate. In blue, 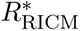, measured using RICM with optical sectioning.The orange and green boxes, 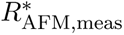 and 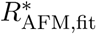, respectively correspond to AFM measurements and fits using equation 7. The bottom images (10.81 *µ*m 10.81 *µ*m) were obtained with RICM and illustrate the distribution of RBC adhesion profiles for each *c*_PLL_ concentration. b) Estimation of the coefficient of variation of membrane fluctuations, *CV* = 100(〈*δI〉/〈I〉 - 〈δI〉/〈I*〉 _background_) with varying *c*_PLL_ concentrations on *N* = 519 cells. Below, stack-normalized RBC fluctuations are displayed for each concentration for gray values [0, 255], while stack-summed RBC images display interference fringes localized at the RBC’s upper membrane.

Furthermore, the geometry of the cell-substrate contact is dependent on PLL concentration. On the one hand, its area *A*^∗^, convexity and ellipticity increase with adhesion and stabilize around *c*_PLL_ = 10^−1^ mg*/*mL, see Figures 2 (a) and S7, suggesting an increase of membrane tension. This is correlated with the decreasing thickness of the white ring that surrounds the cell (Figure 2 (a)), which is a proxy for the contact angle of the cell (the broader the white rim, the smaller the contact angle). In the case of very strong adhesion, the RBC follows the Young-Dupré equation of a wetting droplet, and the contact angle therefore measures the tension of the membrane [34]. All of these observations indicate that the cell’s excess area decreases with increasing PLL concentration.

RICM also provides information on the upper cell membrane, see Figure 1 (c). While the variation coefficient of membrane fluctuations decreases from ≈ 0.5% down to 0% with adhesion (see Figure 2 (b)), the RBC takes an increasingly tensed dome shape, as shown by circular concentric interference rings, see Figure 2 (b). The number of interference rings increases with concentration and stabilize at *c*_PLL_ = 10 mg*/*mL, yielding non-fluctuating domes of height *h*_RBC_ = 2.4 ± 0.2 *µ*m in adequation with the study of stronly-adherent RBCs by Hategan *et al.* and confirmed by confocal microscopy (Figure S6, supplemental materials). In conclusion, while the adhesion cross-section already stabilizes at lower PLL concentrations, the upper shape of the RBC changes progressively into a spherical dome upon increasing PLL coverage. This is unlikely due to pressurisation of the cell since experiments are carried-out in isotonic conditions, but rather the outcome of the competition of adhesion with bending, stretch and shear energies of the membrane [48].

The shape of the upper membrane is further characterized using AFM contact imaging, using the model-adjusted contact point of force curves to evaluate the cell height. Figure 3 (a) shows a radially-averaged 1D profile of a RBC in blue, and the resulting indentation depth for *F* = 30 pN in dark green, in the case of minimal and maximal PLL plate coverage.

**Figure 3:**
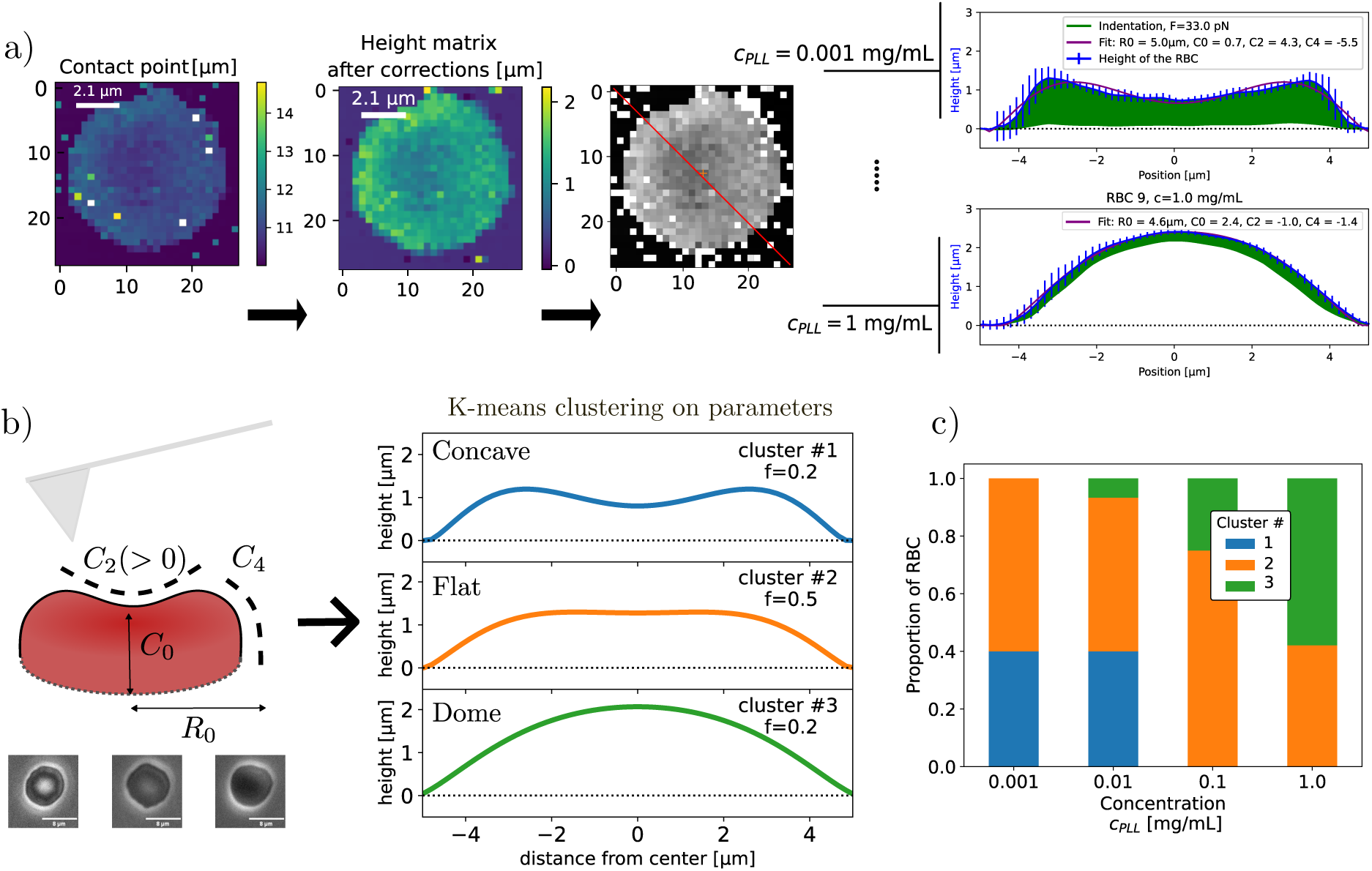
Top surface description of the RBC: a) Height topographies of the RBCs obtained using QI™Advanced Imaging mode. Segmentation and radial averaging is performed to provide the average profile of the RBC. Profile lines of a very soft RBC (*c*_PLL_ = 10^−3^ mg*/*mL) and a dome-shaped RBC (*c*_PLL_ = 1 mg*/*mL) are provided as examples of output of the analysis pipeline. b) Left, schematic description of the RBC top surface using parameters *R*_0_, *C*_0_, *C*_2_ and *C*_4_, as featured in eq. 7. Right, clusters of RBC shapes obtained with unsupervised k-means clustering of geometric parameters with *n* = 3 classes, and their relative frequency in the pool of measured RBC. The silhouette score measuring the cohesion and separation of clusters equals 0.44, which is significant regarding the inherent biological variance. c) Proportions of clusters obtained for each *c*_PLL_ concentration.

This 1D profile is fitted with an equation provided by Fung *et al.* [49] for each PLL concentration (Table S4), although other geometrical models could be relevant as well [50].

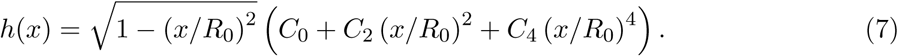

A k-mean unsupervised clustering performed with the set of four variable geometrical parameters *R*_0_*, C*_0_*, C*_2_*, C*_4_ provides 3 classes of typical RBC shapes corroborating RICM observations: a concave, a flat and a dome-shaped RBC, represented on the central panel of Figure 3 (b). Although phase-contrast images of RBCs also seem to cluster into 3 categories featuring clear, gray and dark centers (see Figure 3 (b)), the silhouette score of clustering *S* = 0.34 obtained when mapping the clusters with phase-contrast images is lower than the score *S* = 0.46 obtained when mapping the clusters with PLL concentrations. It means that our study does not permit to sort RBCs to a specific shape cluster using phase contrast during AFM measurements, or at least it is not more efficient than just tuning *c*_PLL_. The respective percentages of clusters for each PLL concentration are displayed on Figure 3 (c), which shows that there is a distribution of the 3 typical shapes for each PLL concentration. The intermediate flat shape is present at each concentration and peaks at *c*_PLL_ = 10^−1^ mg*/*mL, while the concave and dome shapes are most frequent respectively at lowest and highest PLL plate coverage. Analyzing the force-curves measured at the center of the RBCs yields a threefold increase in the corrected Young’s Modulus *E_c_* for dome-shaped RBCs (cluster #3) compared to the concave and flat RBCs (cluster #1 and #2), as shown on Figure S8. It means that concave and flat RBCs are expected to be found with similar elastic and viscoelastic properties, contrary to dome-shape RBCs that are expected to be stiffer. Table S5, Supplemental Materials, provides the average geometric parameters obtained for each cluster. The concave shape is in adequation with Fung *et al.* results for isotonic non-adherent RBCs, although *C*_2_ differs due to flattening. However, a systematic over-estimation of *R*_0_ ≡ *R*^∗^ with the model is observed, compared to the radius *R*^∗^ found with RICM or AFM, see Figure 3.

Since the RBC shape at maximal adhesion follows the Young-Dupré law, it is expected that *V* and *A* remain constant with change of adhesion [47]. However, the height maps obtained with AFM suggest an increasing volume *V* with adhesion, see Table S6. This increased volume is not a consequence of RBC pressurization, since experiments are performed in isotonic conditions. Instead, while most RBCs height maps are performed on glutaraldehyde-fixed cells to avoid the pitfall of the extreme softness of the RBC, imaging live RBCs in contact mode is performed at the cost of very large indentations spanning the whole RBC. As a consequence, the contact-point is ill-defined, as observed by a maximal RBC thickness which is lower than obtained with optical methods. Supplemental Materials (Table S6) provide an estimation of the contact point error by calculating the volume difference with the typical volume of a healthy adult RBC.

In summary, this study provides a glimpse into the continuum of shapes driven by non-specific adhesion to the substrate, which is described theoretically in the case of vesicles but seldom observed experimentally: RBCs undergo impressive shape modifications with substrate adhesion, even for very low concentrations of plate coating. Those shapes can be classified into concave, flat and dome-shaped RBC, however cannot be simply isolated with concentration tuning. Furthermore, recent results [51] suggest that PLL coverage could actively modulate the shape and mechanics in other types of cells: CaF_2_ PLL-coated surfaces with different incubation times show loss of cell polarisation and modified mechanical properties, accompanied by an increase of apoptotic markers and modifications of the secondary structure of proteins.

### The rigidity of the RBC increases with substrate adhesion

To correlate the RBC’s geometry with its mechanical properties upon non-specific substrate adhesion, a parametric study of the Young’s modulus *E* measured with AFM as a function of PLL concentration was performed over a larger dataset involving 11 different donors. In order to reduce the indentation depth and provide a single non-local measurement of stiffness for each RBC, indentation is performed using a 3 *µ*m diameter spherical bead, in contrast with the 70 nm radius probe used for height maps. In Figure 4 (a) are plotted violin plots showing the RBcs’ stiffness analyzed with -*E_c_*- and without -*E*- substrate correction. *E_c_*decreases by an order of magnitude relative to *E*, stemming from the large indentations obtained for the setpoint *F* = 30 pN (Figure S4 b, Supplemental Materials), which nevertheless represents the lowest accessible setpoint ensuring a reasonable signal to noise ratio. *E_c_* depends significantly on concentration *c*_PLL_, even though the correction factor *g*(*χ*_0_)^−1^ decreases monotonically with *c*_PLL_ (Figure S4 c, Supplemental Materials), which narrows the gap between the values of *E_c_*at weak and strong substrate adhesion. Statistical analysis of the *E_c_* values is available in Table 2: it appears that *E_c_* ∈ [10 − 100] Pa, which is in the lowest range reported in the literature (see Table S8).

**Figure 4:**
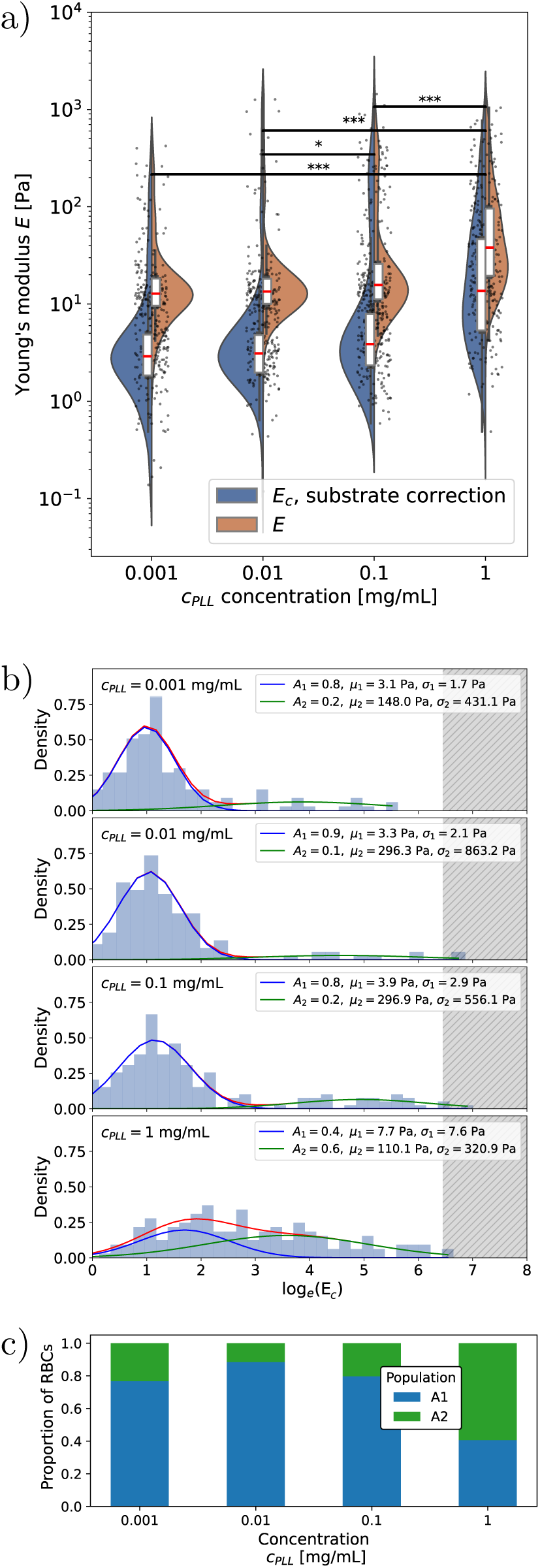
a) Young’s modulus distribution for *F*_setpoint_ = 30 pN for all *c*_PLL_ concentration, without (orange box) and with (blue box) substrate corrections for the rigid substrate effect. b) Distribution of the corrected Young’s Moduli for *F*_setpoint_ = 30 pN, fitted with 2 lognorm distributions for each concentration in blue and in green respectively, where *A_i_* is the population frequency, *µ_i_*the mean of the distribution in Pa, and *σ_i_* its standard deviation in Pa. The red curve displays the cumulative distribution. c) Bar chart of the stacked proportions of populations 1 (blue) and 2 (green) as a function of *c*_PLL_.

**Table 2:** Young’s modulus *E* of RBCs for various PLL incubation concentrations, after correction of the effect of the underlying rigid substrate [42].

| $c_{\text{PLL}}$ [mg/mL] | $E$ (mean) [Pa] | s.e.m. [Pa] | $E$ (median) [Pa] | Q1 [Pa] | Q3 [Pa] |
| --- | --- | --- | --- | --- | --- |
| $10^{-3}$ | 12.41 | 2.90 | 2.99 | 1.84 | 4.87 |
| $10^{-2}$ | 26.37 | 10.06 | 3.11 | 1.97 | 4.81 |
| $10^{-1}$ | 41.92 | 9.18 | 3.88 | 2.34 | 7.95 |
| $10^0$ | 59.32 | 9.73 | 13.69 | 5.36 | 46.42 |

In Figure 4 (b), the continuum of values found for *E* is indeed well described with a finite number of log-normal distributions. We find that a single distribution centered on *µ*_1_ ≈ 15 Pa without substrate correction and *µ*_1_ ≈ 3 Pa with substrate correction is retrieved for each concentration. The associated population *A*_1_ decreases with PLL coverage, while standard deviation *σ*_1_ increases. Similarly, a second wider distribution is found for each concentration, which population *A*_2_ increases with PLL coverage. We attributed the first population to the pool of concave and flat RBCs, and the second population to dome-shaped RBCs. The population frequencies *A*_1_ and *A*_2_ shown in Figure 4 (c) are in line with the clusters found in Figure 3 (c), and it corroborates the observation that dome-shaped RBCs have a wider and higher distribution of effective moduli (Figure S8). We also observe that *E_c,_*_max_ ≈ 1 kPa is never exceeded for each *c*_PLL_ concentration, and might correspond to the maximum value before lysis occurs (hatched in gray on Figure 4 (b)). Experimentally, the frequency of spontaneous lysis is observed to increase with *c*_PLL_, due to the number of RBCs that are already tensed by adhesion. It stresses the importance of performing experiments within the spontaneous RBC lysis time *t* ≈ 40 min [34] which occurs when area stretching exceeds 3% [8].

In summary, substrate adhesion affects the mechanical properties of RBCs measured with AFM. It appears that the set of shapes triggered by PLL also correlate with defined distributions of Young’s moduli, the dome-shape RBCs forming a population with higher Young’s Moduli.

### A clear limitation of the Hertz model

Consistent with the previous section, mechanical properties of RBCs have been extensively characterized by their effective Young’s modulus *E* using AFM, adjusted to force-indentation curves under the assumption of an isotropic, elastic, semi infinite flat sample. However, Figure S9 (a) and (b) (Supplemental Materials) shows that the values for *E* approached by the Hertz model following *F* (*δ*) ∝ *δ*^3^*^/^*^2^ depend on the maximal force used to fit the model. Curves for 5 RBCs adhered at different *c*_PLL_ concentrations from 10^−3^ mg*/*mL to 10^1^ mg*/*mL are displayed, where (a) and (b) respectively correspond to the adjusted model for *F* = 30 pN and *F* = 1 nN. These setpoints, respectively, correspond to the force used in our experiments, and the maximal force range used in the literature. Strikingly, the determination of the contact point and the magnitude of *E* are severely dependent on *F* : in the case of 1% coverslip coverage by PLL - a very common substrate treatment (*c*_PLL_ = 0.001 mg*/*mL) - *E* varies from *E* ≈ 10 Pa at *F* = 30 pN to *E* ≈ 10^4^ Pa at *F* = 1 nN. Table S8 in the Supplemental Materials provides a detailed literature survey corroborating that higher *F* values correlate with higher *E* values in other studies as well. However, coating concentrations are not consistently reported, which does not permit comparison with our study.

However, it appears that the adequation of the Hertz model decreases when the force setpoint increases. To investigate this observation, Figure S9 (c) shows, using Hertz model for *F* ≤ 200 pN across a large pool of RBCs, that increasing the force setpoint *F* leads to higher *E* for all concentrations, with a pronounced effect at lower *c*_PLL_ values. A direct explanation is found by adjusting a power-law *F* ∝ *δ^b^*with exponent *b* and free contact point as a function of force setpoint *F*, see Figure S9 (d). For *F* ≈ 30 pN, all adjustments collapse in an exponent *b* ≈ 1.5 in line with Hertz model. Conversely, increasing *F* results in non-physical exponents for concentrations *c*_PLL_ ≤ 10^−1^ mg*/*mL, most likely due to a strong substrate effect: usual corrections introduced previously [42] suppose an indentation small enough that the exponent *b* remains unchanged, which reveals insufficient for high force setpoints. For concentrations *c*_PLL_ ≥ 1 mg*/*mL, *b* increases before stalling around *b* ≈ 3. Therefore, only very low force setpoints, as used in this study, lay in the suitable range for using the Hertz model. However, for strong cell-substrate adhesion a stable regime is found at higher setpoints, suggesting that another model than Hertz would fit the data better.

Hence, studying RBCs using the Hertz model shows clear limits: not only do cells undergo large deformations and therefore the underlying rigid substrate becomes non-negligible, but the hypothesis of an isotropic elastic sample also becomes unjustified. This is where the different shape regimes of RBCs depending on PLL concentration come into play. As already reported by Hategan *et al.* [34], we observe similarities in the shapes observed by RBCs under adhesion and the theoretical predictions for deflated skeleton-free vesicles obtained by minimising the membrane free-energy in the Helfrisch-Canham formalism [48]. In the strong adhesion regime - which occurs here for *c*_PLL_ ≥ 1 mg*/*mL - the RBC takes a spherical cap shape, as predicted for vesicles when surface tension becomes the main competitive term with adhesion instead of bending. In this regime, Hategan *et al.* [34] and Sen *et al.* [52] proposed an analytical and a computational membrane indentation model which is a linear combination of a linear term *F* (*δ*) ∝ *δ* accounting for pre-stress *T*_0_ of the membrane, and a cubic term *F* (*δ*) ∝ *δ*^3^ accounting for membrane area dilation with coefficient *K*. They show that the linear term is dominant over the first 100 − 200 nm of indentation. This model corroborates the behaviour of exponent 2 *< b <* 4 for *c*_PLL_ ≥ 1 mg*/*mL.

Here, however, we also span lower adhesion regimes where membrane tension is not expected to be dominant compared to bending, especially at *c*_PLL_ = 10^−3^ mg*/*mL where the concave shaped RBCs represent a large fraction of the population. Barns *et al.* conducted force curve simulations including bending, stretching and area dilation contributions to RBC deformation, and established the dominance of bending over other deformation modes for dome-shaped RBCs [53]. Similarly to prestress, bending also implies a linear increase of F with indentation: *F* (*δ*) ∝ *δ* [54].

For that purpose, we propose a simple continuous model assuming non-coexistent bending and area stretching, which is adjusted for *F*_setpoint_ ≤ 200 pN on Figure 5 (a), using the accurate contact-point estimated with *F*_setpoint_ = 30 pN (*δ* = 0).

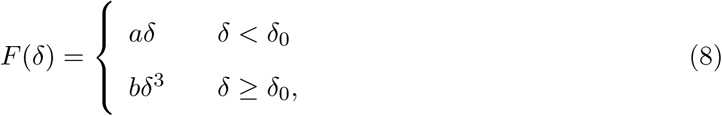

**Figure 5:**
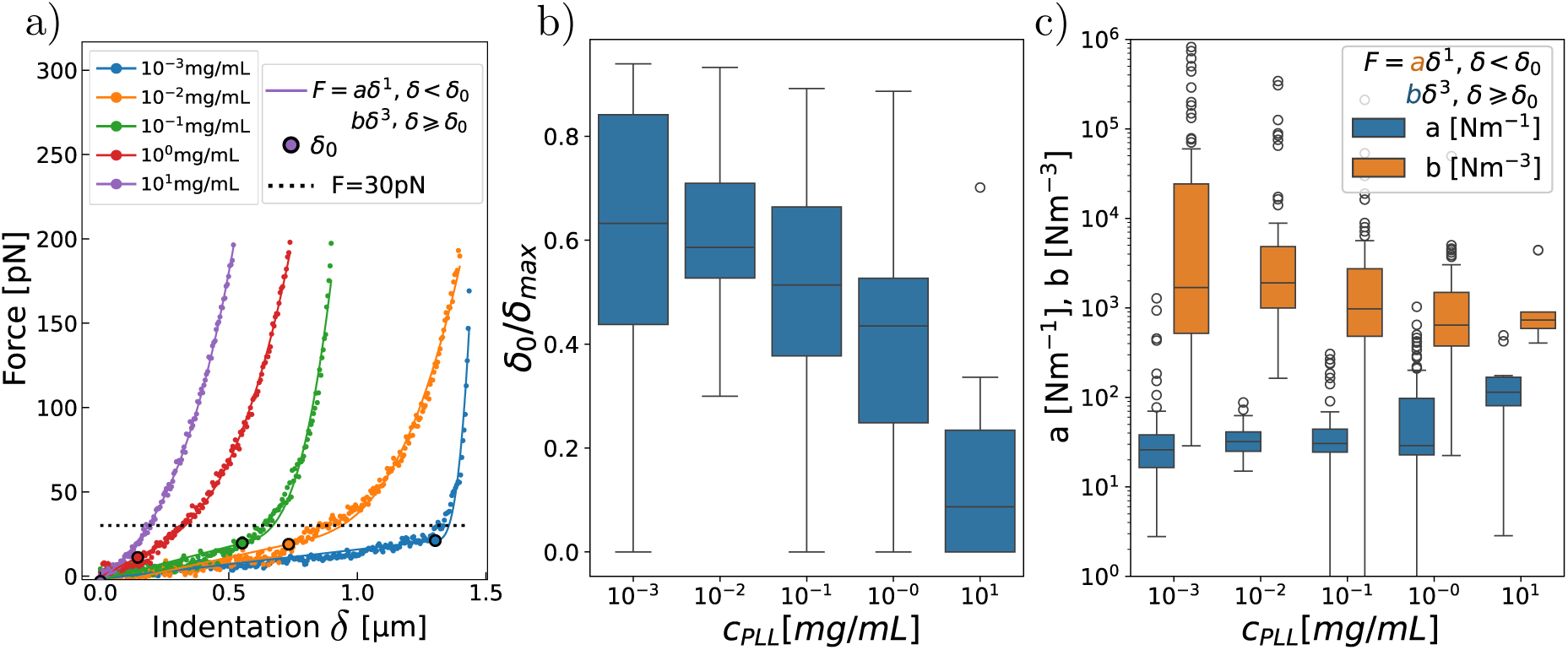
*F*_setpoint_ = 200 pN. a) Piecewise power law model (Eq 8, 2 free parameters *a* and *b* fitted on 5 typical force-indentation curves for *c*_PLL_ [10^−3^, 10^1^] mg*/*mL. *δ*_0_, the indentation value indicating transition from linear (bending dominated) to cubic (stretching dominated) power-law, is represented as a dot marker outlined in black. b) Distribution of the bending-dominated indentation ratio *δ*_0_*/δ*_max_, where *δ*_max_ is calculated for *F*_setpoint_ = 200 pN, over *N* = 306 cells. c) Distribution of prefactors *a* and *b*, respectively accounting for the linear and cubic contributions to force-indentation curves.

with *a* and *b* constants. The goodness of fit surpasses that of Hertz model, and provides an approximate distance *δ*_0_ separating a bending dominated deformation from an area stretching deformation. Figure 5 (b) shows the tendency of *δ*_0_*/δ*_max_ to decrease with *c*_PLL_, since stretching becomes the dominant mode of deformation at strong adherence. On Figure 5, *a* and *b* are the prefactors to the linear and cubic terms. *a* is constant in the low adhesion regime (*c*_PLL_ ≤ 0.1 mg*/*mL) and provides an indirect measurement of bending. It increases for stronger adhesion where it reflects the pre-stress of the membrane. *b* provides an indirect measurement of the membrane elastic stretching constant for *c*_PLL_ ≥ 1 mg*/*mL. However, the broad and elevated distribution of *b* for low *c*_PLL_ reflects the influence of the substrate, which cannot be corrected for such soft samples since the assumption of small deformations breaks down for *F* = 200 pN [42]. Therefore, we suggest that with increasing PLL concentration, the effective Young’s modulus *E* measured with AFM is affected by the decrease of excess area, and area stretching becomes the major mode of deformation : this is indeed how *K*, the expansion coefficient, is measured in micropipette experiments [12]. Hence, we interpret the bimodal distribution of Young’s moduli (Figure 4) as the signature of different modes of deformation being probed during indentation, depending on the strength of adhesion, due to transition from a deformation driven by bending (weak adhesion) to a deformation driven by membrane stretching (strong adhesion). Consequently, in order to properly investigate the mechanics of RBCs with AFM, it could be necessary to develop a non-linear indentation model combining the different deformation modes of the RBC, similarly to what has been performed for vesicles [55] or epithelial sheets of cells [56]. Such a model could make sense of the values found for the prefactors of our phenomenological model, Equation 8. However, other formalisms are required than previously proposed for nanovesicles, which are pressurized [57], or GUVs, for which bending is negligible [55].

### The rheology of the RBC is impacted by the adhesion strength to the substrate

To further understand the role of non-specific adhesion in the measured mechanical properties of RBCs, viscoelasticity is probed with AFM by applying a sinusoidal deformation of constant amplitude, small with respect to indentation *δ*_0_, using a force setpoint of *F* = 30 pN in order to remain in the regime where Hertz model can be applied. The sinusoidal force response measured with the cantilever deflection provides a frequency-dependent measure of the complex shear modulus *G*^∗^(*ω*). The model introduced for the study of the shear modulus is the empirical power law structural damping model as described in Fabry *et al.* [58],

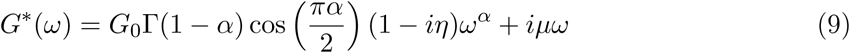

Where *ω* = 2*πf* is the angular frequency of the probe, *G*^∗^ follows a power law with exponent *α* and accounts for an additional Newtonian viscosity *µ* that becomes significant with increasing *f* values. 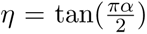 is the hysteresis term, which denotes the coupling between the loss and shear moduli. The choice of the model is guided by the linear behaviour of *G*′′(*f*) for *f >* 10 Hz, while *G*′(*f*) deviates from a plateau for lower PLL concentrations. The model captures better both the shear and loss moduli compared to a simple Kelvin Voigt model, as shown by *α* ≠ 0. A comparison is provided in Figure S10, Supplemental Materials. The loss and shear moduli are respectively the imaginary and real parts of *G*^∗^, defining the dissipated and stored components of the isotropic shear modulus. The loss tangent *G*′′*/G*′, which quantifies at which frequency the system dissipates more energy (flows) than it stores, defines a cut-off frequency *f_t_*when *G*′ = *G*′′. The shear and loss moduli are plotted on Figure 6 (a) as open round and filled triangle markers, respectively, and represented as a function of frequency *f*.

**Figure 6:**
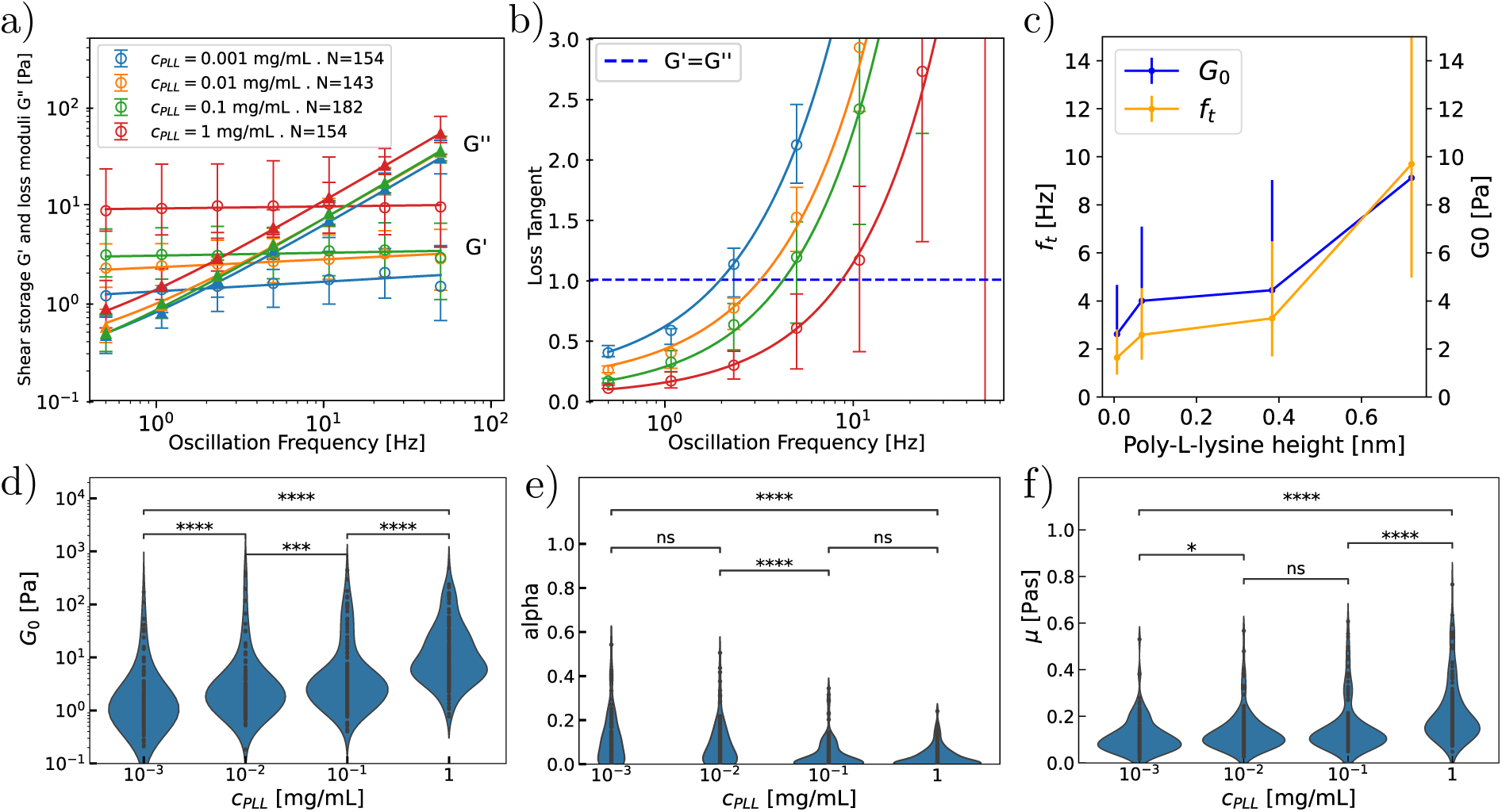
a) Corrected shear modulus *G*′ and loss modulus *G*′′ of single RBCs for varying *c*_PLL_ concentrations. Open round markers correspond to *G*′, filled triangle markers to *G*′′. *N* corresponds to the number of cells per condition. b) Corresponding loss tangent *G*′′*/G*′. In a) and b), error bars represent the median interquartile range (Q1–Q3), and the full lines correspond to the structural damping model (eq. 9) conjointly adjusted for *G*′ and *G*′′. c) *f_t_* (blue curve) and *G*_0_ (orange curve) are plotted as a function of PLL height *h*_PLL_ (eq. 1) which is proportional to the effectively adsorbed concentration of PLL. Parameters are obtained by minimizing the least-square cost function. Error bars represent the median ± interquartile range (Q1–Q3). d -f), Distribution of fit parameters *G*_0_, *α* and *µ* (respectively) for each *c*_PLL_ concentration.

The frequency dependence of the complex shear modulus reveals the rheological properties of RBCs in different adhesion regimes: the loss modulus *G*′′(*f*) is almost not modified by the PLL surface coverage, indicating that the viscous properties of the RBCs are left unchanged with adhesion, except for *c*_PLL_ = 1 mg*/*mL in the strong adhesion regime. On the other hand, the shear modulus *G*_0_ increases by one order of magnitude with *c*_PLL_, implying that the frequency *f_t_* increases with *c*_PLL_ (Figure 6 (b)). This is shown on Figure 6 (c), where *G*_0_ and *f_t_* are displayed in blue and orange respectively. Using equation 1, they are plotted against *h*_PLL_, which measures the adsorbed concentration of PLL. It appears that both *f_t_* and *G*_0_ follow a comparable linear trend with *h*_PLL_, since the modification of *G*′(*f*) is primarily responsible for the change in viscoelasticity of the RBC: the cell behaves like a solid on a larger range of frequencies for strong adhesion.

Figure 6 (e-g) displays the distributions of the three fitting parameters *G*_0_, *α* and *µ* (respectively). Their mean values and standard error for each concentration *c*_PLL_ are reported in Table 3. *G*_0_ measured at *f* = 0.5 Hz verifies approximately *E_c_*= 2*G*_0_(1 + *ν*) with *ν* = 0.5, and is in the lowest range found in the literature, which might arise from the use of a very low setpoint force: it is known that RBCs show an apparent stiffening with increasing applied forces [26], corroborated by the findings of the previous section. Viscosity *µ* is in line with the order of magnitudes obtained for the membrane’s viscosity in other RBC viscoelastic studies, using optical tweezers (Betz *et al.*, *η* = 6.4 ± 1.9 Pas [4]) or dRT-DC (Fregin *et al.* [59] find a Kelvin-Voigt viscoelastic behavior of single RBCs with *G*_0_ = 100 ± 10 Pa and *η* = 0.24 ± 0.18 Pa.s). Concerning *α*, observations show that it decreases continuously to 0 with increasing cell-substrate adhesion. This means that with increasing adhesion, the RBC viscoelastic properties tend towards a simple Kelvin-Voigt model describing a viscoelastic solid. This is in contradiction with former studies described in Table S7, which obtained an exponent *n* in the range [0.5, 0.7] for the loss modulus using a power law *G*′′(*f*) ∝ *f^n^*, while we obtain *n* → 1 with increasing *c*_PLL_.

**Table 3:** Table of fitting parameters obtained for different *c*_PLL_ concentrations, introducing the mean and STD for each parameter.

| | $G_0$ [Pa] | | $\alpha$ | | $\mu$ [Pa.s] | | $f_t$ [Hz] | |
| --- | --- | --- | --- | --- | --- | --- | --- | --- |
| PLL Concentration [mg/mL] | mean | s.e. | mean | s.e. | mean | s.e. | mean | s.e. |
| $10^{-3}$ | 4.68 | 1.4 | 0.11 | 0.01 | 0.11 | 0.01 | 4.67 | 0.01 |
| $10^{-2}$ | 11.4 | 4 | 0.09 | 0.01 | 0.13 | 0.01 | 6.96 | 0.93 |
| $10^{-1}$ | 15.9 | 3.41 | 0.04 | 0.00 | 0.15 | 0.01 | 8.78 | 0.88 |
| 1 | 28.1 | 4.5 | 0.03 | 0.00 | 0.21 | 0.01 | 15.1 | 1.15 |

The analysis of a set of *N* = 8 comparable blood donors confirms a robust trend in the significant change in parameters from low to high adhesion of RBCs, see Figure S11. The strong variability between donors can stem from inter-donor variability, but also variability of PLL coating uniformity or different samplings of RBC ages, and does not permit more in-depth analysis of this dataset.

The work presented here provides a microrheology study of single RBCs, raising comparable results to other methods used at the single-cell level [24, 60]. It shows that substrate-adhesion impacts the frequency-dependent response of the RBC, mostly through the elastic component *G*′(*f*) of the complex shear modulus. Fedosov *et al.* [61, 62] also found a linear increase of *G*′′(*f*) with *f* and a drastic increase of *G*′(*f*) when modelling pathology-mediated cytoskeleton stiffening, which stems from their assumption of an increasing membrane shear modulus without change in cytosolic or membrane viscosity. Likewise, they obtain a transition frequency *f_t_*≈ 4 Hz for healthy RBCs and *f_t_* ≈ 13 Hz for last-stage malaria-infected RBCs. Besides pathology or adhesion strength, structural modifications following changes in osmolarity are also known to alter the measured mechanical properties of RBCs: Park *et al.* [17] experimentally showed that bending rigidity, area modulus and shear modulus increase with external osmolarity, while the height of membrane fluctuations decreases. We also suggest that different levels of ATP-mediated activity of RBCs depending on adhesion strength could explain that *G*′(*f*) follows different trends at low frequency. Indeed, the active transient detachment of spectrin from the lipid membrane provides a dynamic softening of the membrane and accounts for the non-equilibrium dynamics of the membrane for *f <* 10 Hz [4, 5]. Out-of-equilibrium dynamics might vanish in the strong adhesion regime, since membrane fluctuations decay.

## Conclusion

Non-specific substrate adhesion is key in immobilizing cells in many in-vitro experiments. The present study, which uses RICM and AFM to elucidate both the shape and rheology of RBCs in different adhesion regimes, shows that non-specific substrate adhesion of RBCs mediated by Poly-L-Lysine in the incubation concentration range 10^−3^ −1 mg*/*mL triggers a transition in the shape of RBCs, from a biconcave to a dome shape. This transition impacts the mechanical properties of RBCs measured with AFM: it is accompanied by an increase in the stiffness *E* of the RBCs, which follows a bimodal log-norm distribution that discriminates strongly attached dome-shaped RBCs from concave or flat RBCs. Furthermore, the rheology of RBC is evaluated on the frequency range *f* ∈ [0.5, 100] Hz. While the storage modulus *G*′(*f*) increases strongly with PLL concentration, the loss modulus *G*′′(*f*) is left unchanged even in the strong adhesion regime. This implies that the transition frequency *f_t_*, separating the dominantly elastic and viscous regimes, shifts with adhesion. Furthermore, standard contact mechanics models assuming isotropic materials, although successfully applied to other types of cells, fail to describe the indentation of RBCs. It stems from the leading role of the 2D membrane in the RBCs deformation, as well as its extreme deformability. We show that the Hertz model is valid only for indentation forces below *F* = 30 pN: above that range, the stiffness of weakly attached RBCs, bending-dominated, is largely overestimated by the influence of the rigid substrate. Conversely, the stiffness of strongly attached RBCs is dominated by the stretching of the tensed membrane. Furthermore, the adhesion of RBCs is mostly triggered by chemical affinity of the membrane with the surface, and does not involve active protein complexes. Our findings imply that in-vitro measurements of substrate-adherent cells, in particular RBCs, require careful control of adhesion to ensure reproducibility. In addition to adhesion strength, the wide range of rigidification treatments used (formaldehyde, glutaraldehyde), different tip geometries and approach parameters, as well as data-processing and contact-point identification, explain the broad distribution in effective Young’s moduli found in the literature.

## Author Contributions

A.N., C.V. and D.D. conceived the experiments, A.N. and D.D. conducted the experiments, A.N. analyzed the results. C.V. conducted the project. A.N. wrote the initial draft, all authors reviewed the manuscript.

## Acknowledgements

Authors acknowledge the support of the French Agence Nationale de la Recherche (ANR) with grant ANR-21-CE09-0011 (CellDance project), and the help of Lionel Bureau for ellipsometry experiments and the preparation of silicon wafers. Authors also acknowledge the help of Taha Benyattou in the administration of the CellDance project, and Alexander Erlich, Magalie Faivre, Lotfi Berguiga and Ha My Dang Nguyen for fruitful discussions.

## Data availability

Data and python codes are available upon reasonable request.

## Competing interests

The authors declare no competing interests.

## Notes

### Competing Interest Statement

The authors have declared no competing interest.

## References

[1] Alexis Moreau, François Yaya, Huijie Lu, Anagha Surendranath, Anne Charrier, Benoit Dehapiot, Emmanuèle Helfer, Annie Viallat, and Zhangli Peng. Physical mechanisms of red blood cell splenic filtration. Proceedings of the National Academy of Sciences, 120(44): e2300095120, October 2023. ISSN 0027-8424, 1091-6490. doi: 10.1073/pnas.2300095120.

[2] S. Chien, R. G. King, R. Skalak, S. Usami, and A. L. Copley. Viscoelastic properties of human blood and red cell suspensions. Biorheology, 12(6):341–346, October 1975. ISSN 0006-355X. doi: 10.3233/bir-1975-12603.

[3] Stuart M Cahalan, Viktor Lukacs, Sanjeev S Ranade, Shu Chien, Michael Bandell, and Ardem Patapoutian. Piezo1 links mechanical forces to red blood cell volume. eLife, 4: e07370, May 2015. ISSN 2050-084X. doi: 10.7554/eLife.07370.

[4] Timo Betz, Martin Lenz, Jean-François Joanny, and Cécile Sykes. ATP-dependent mechanics of red blood cells. Proceedings of the National Academy of Sciences, 106(36):15320–15325, September 2009. doi: 10.1073/pnas.0904614106.

[5] H. Turlier, D. A. Fedosov, B. Audoly, T. Auth, N. S. Gov, C. Sykes, J.-F. Joanny, G. Gompper, and T. Betz. Equilibrium physics breakdown reveals the active nature of red blood cell flickering. Nature Physics, 12(5):513–519, May 2016. ISSN 1745-2481. doi: 10.1038/nphys3621.

[6] P. B. Canham. The minimum energy of bending as a possible explanation of the biconcave shape of the human red blood cell. Journal of Theoretical Biology, 26(1):61–81, January 1970. ISSN 0022-5193. doi: 10.1016/S0022-5193(70)80032-7.

[7] W. Helfrich. Elastic Properties of Lipid Bilayers: Theory and Possible Experiments. Zeitschrift für Naturforschung C, 28(11-12):693–703, December 1973. ISSN 1865-7125. doi: 10.1515/znc-1973-11-1209.

[8] E. A. Evans and R. Skalak. Mechanics and thermodynamics of biomembranes: Part 1. CRC critical reviews in bioengineering, 3(3):181–330, October 1979. ISSN 0045-642X.

[9] E. A. Evans. Bending elastic modulus of red blood cell membrane derived from buckling instability in micropipet aspiration tests. Biophysical Journal, 43(1):27–30, July 1983. ISSN 0006-3495. doi: 10.1016/S0006-3495(83)84319-7.

[10] R M Hochmuth, P R Worthy, and E A Evans. Red cell extensional recovery and the determination of membrane viscosity. Biophysical Journal, 26(1):101–114, April 1979. ISSN 0006-3495. doi: 10.1016/S0006-3495(79)85238-8.

[11] E. A. Evans and R. Waugh. Osmotic correction to elastic area compressibility measurements on red cell membrane. Biophysical Journal, 20(3):307–313, December 1977. ISSN 0006-3495. doi: 10.1016/S0006-3495(77)85551-3.

[12] E. A. Evans. Structure and deformation properties of red blood cells: Concepts and quantitative methods. Methods in Enzymology, 173:3–35, 1989. ISSN 0076-6879. doi: 10.1016/s0076-6879(89)73003-2.

[13] A. Katchalsky, O. Kedem, C. Klibansky, and A. DeVries. Rheological considerations of the haemolysing red blood cell. Flow Properties of Blood and Other Biological Systems, page 155–171, December 1976. ISSN 9780124019508. doi: 10.1016/B978-0-12-401950-8.50032-0.

[14] T. W. Hoeber and R. M. Hochmuth. Measurement of red cell modulus of elasticity by in-vitro and model cell experiments. Journal of Basic Engineering, 92(3):604–609, 1970. ISSN 0021-9223. doi: 10.1115/1.3425084.

[15] R.M. Hochmuth, N. Mohandas, and P.L. Blackshear. Measurement of the elastic modulus for red cell membrane using a fluid mechanical technique. Biophysical Journal, 13(8):747–762, August 1973. ISSN 00063495. doi: 10.1016/S0006-3495(73)86021-7.

[16] I. M. Lamzin and R. M. Khayrullin. The quality assessment of stored red blood cells probed using atomic-force microscopy. Anatomy Research International, 2014(1):869683, 2014. ISSN 2090-2751. doi: 10.1155/2014/869683.

[17] YongKeun Park, Catherine A. Best, Kamran Badizadegan, Ramachandra R. Dasari, Michael S. Feld, Tatiana Kuriabova, Mark L. Henle, Alex J. Levine, and Gabriel Popescu. Measurement of red blood cell mechanics during morphological changes. Proceedings of the National Academy of Sciences of the United States of America, 107(15):6731–6736, April 2010. ISSN 0027-8424. doi: 10.1073/pnas.0909533107.

[18] Ida Dulińska, Marta Targosz, Wojciech Strojny, Małgorzata Lekka, Paweł Czuba, Walentyna Balwierz, and Marek Szymoński. Stiffness of normal and pathological erythrocytes studied by means of atomic force microscopy. Journal of Biochemical and Biophysical Methods, 66 (1-3):1–11, March 2006. ISSN 0165022X. doi: 10.1016/j.jbbm.2005.11.003.

[19] Sylvie Hénon, Guillaume Lenormand, Alain Richert, and François Gallet. A New Determination of the Shear Modulus of the Human Erythrocyte Membrane Using Optical Tweezers. Biophysical Journal, 76(2):1145–1151, February 1999. ISSN 0006-3495. doi: 10.1016/S0006-3495(99)77279-6.

[20] Georgii V. Grigorev, Alexander V. Lebedev, Xiaohao Wang, Xiang Qian, George V. Maksimov, and Liwei Lin. Advances in Microfluidics for Single Red Blood Cell Analysis. Biosensors, 13(1):117, January 2023. ISSN 2079-6374. doi: 10.3390/bios13010117.

[21] Sebastian Himbert, Angelo D’Alessandro, Syed M. Qadri, Michael J. Majcher, Todd Hoare, William P. Sheffield, Michihiro Nagao, John F. Nagle, and Maikel C. Rheinstädter. The bending rigidity of the red blood cell cytoplasmic membrane. PLOS ONE, 17(8):e0269619, August 2022. ISSN 1932-6203. doi: 10.1371/journal.pone.0269619.

[22] M. Shahrooz Amin, YougKeun Park, Niyom Lue, Ramachandra R. Dasari, Kamran Badizadegan, Michael S. Feld, and Gabriel Popescu. Microrheology of red blood cell membranes using dynamic scattering microscopy. Optics Express, 15(25):17001, 2007. ISSN 1094-4087. doi: 10.1364/OE.15.017001.

[23] Ru Wang, Huafeng Ding, Mustafa Mir, Krishnarao Tangella, and Gabriel Popescu. Effective 3d viscoelasticity of red blood cells measured by diffraction phase microscopy. Biomedical Optics Express, 2(3):485, March 2011. ISSN 2156-7085. doi: 10.1364/BOE.2.000485.

[24] Marina Puig-De-Morales, Mireia Grabulosa, Jordi Alcaraz, Joaquim Mullol, Geoffrey N. Maksym, Jeffrey J. Fredberg, and Daniel Navajas. Measurement of cell microrheology by magnetic twisting cytometry with frequency domain demodulation. Journal of Applied Physiology, 91(3):1152–1159, September 2001. ISSN 8750-7587, 1522-1601. doi: 10.1152/jappl.2001.91.3.1152.

[25] Fran Gómez, Leandro S. Silva, Glauber Ribeiro De Sousa Araújo, Susana Frases, Ana Acacia S. Pinheiro, Ubirajara Agero, Bruno Pontes, and Nathan Bessa Viana. Effect of cell geometry in the evaluation of erythrocyte viscoelastic properties. Physical Review E, 101 (6):062403, 2020. ISSN 2470-0045, 2470-0053. doi: 10.1103/PhysRevE.101.062403.

[26] Giulia Bergamaschi, Kees-Karel H. Taris, Andreas S. Biebricher, Xamanie M. R. Seymonson, Hannes Witt, Erwin J. G. Peterman, and Gijs J. L. Wuite. Viscoelasticity of diverse biological samples quantified by acoustic force microrheology (afmr). Communications Biology, 7 (1):1–14, 2024. ISSN 2399-3642. doi: 10.1038/s42003-024-06367-3.

[27] Yuri M. Efremov, Takaharu Okajima, and Arvind Raman. Measuring viscoelasticity of soft biological samples using atomic force microscopy. Soft Matter, 16(1):64–81, December 2019. ISSN 1744-6848. doi: 10.1039/C9SM01020C.

[28] Shamik Sen, Shyamsundar Subramanian, and Dennis E. Discher. Indentation and Adhesive Probing of a Cell Membrane with AFM: Theoretical Model and Experiments. Biophysical Journal, 89(5):3203–3213, November 2005. ISSN 0006-3495. doi: 10.1529/biophysj.105.063826.

[29] Dina Baier, Torsten Müller, Thomas Mohr, and Ursula Windberger. Red Blood Cell Stiffness and Adhesion Are Species-Specific Properties Strongly Affected by Temperature and Medium Changes in Single Cell Force Spectroscopy. Molecules, 26(9):2771, January 2021. ISSN 1420-3049. doi: 10.3390/molecules26092771.

[30] Mi Li, LianQing Liu, Ning Xi, YueChao Wang, ZaiLi Dong, XiuBin Xiao, and WeiJing Zhang. Atomic force microscopy imaging and mechanical properties measurement of red blood cells and aggressive cancer cells. Science China Life Sciences, 55(11):968–973, November 2012. ISSN 1869-1889. doi: 10.1007/s11427-012-4399-3.

[31] Elena Kozlova, Aleksandr Chernysh, Ekaterina Manchenko, Viktoria Sergunova, and Viktor Moroz. Nonlinear Biomechanical Characteristics of Deep Deformation of Native RBC Membranes in Normal State and under Modifier Action. Scanning, 2018(1):1810585, 2018. ISSN 1932-8745. doi: 10.1155/2018/1810585.

[32] Viktoria Sergunova, Stanislav Leesment, Aleksandr Kozlov, Vladimir Inozemtsev, Polina Platitsina, Snezhanna Lyapunova, Alexander Onufrievich, Vyacheslav Polyakov, and Ekaterina Sherstyukova. Investigation of Red Blood Cells by Atomic Force Microscopy. Sensors (Basel, Switzerland), 22(5):2055, March 2022. ISSN 1424-8220. doi: 10.3390/s22052055.

[33] Alina Hategan, Kheya Sengupta, Samuel Kahn, Erich Sackmann, and Dennis E. Discher. Topographical Pattern Dynamics in Passive Adhesion of Cell Membranes. Biophysical Journal, 87(5):3547–3560, November 2004. ISSN 00063495. doi: 10.1529/biophysj.104.041475.

[34] Alina Hategan, Richard Law, Samuel Kahn, and Dennis E. Discher. Adhesively-Tensed Cell Membranes: Lysis Kinetics and Atomic Force Microscopy Probing. Biophysical Journal, 85 (4):2746–2759, October 2003. ISSN 0006-3495.

[35] Siddhartha Varma, Lionel Bureau, and Delphine Débarre. The conformation of thermoresponsive polymer brushes probed by optical reflectivity. Langmuir, 32(13):3152–3163, April 2016. ISSN 0743-7463, 1520-5827. doi: 10.1021/acs.langmuir.6b00138.

[36] Cathie Ventalon, Oksana Kirichuk, Yotam Navon, Yan Chastagnier, Laurent Heux, Ralf P Richter, Lionel Bureau, and Delphine Débarre. Optical sectioning for reflection interference microscopy: quantitative imaging at soft interfaces. Langmuir, 41(15):10040–10051, 2025.

[37] Cathie Ventalon, Agathe Nidriche, and Delphine Débarre. Quantifying optical sectioning in reflection microscopy with patterned illumination. Biomed. Opt. Express, 17(7):3352–3374, Jul 2026. doi: 10.1364/BOE.593117.

[38] Johannes Schindelin, Ignacio Arganda-Carreras, Erwin Frise, Verena Kaynig, Mark Longair, Tobias Pietzsch, Stephan Preibisch, Curtis Rueden, Stephan Saalfeld, Benjamin Schmid, Jean-Yves Tinevez, Daniel James White, Volker Hartenstein, Kevin Eliceiri, Pavel Tomancak, and Albert Cardona. Fiji: An open-source platform for biological-image analysis. Nature Methods, 9(7):676–682, July 2012. ISSN 1548-7105. doi: 10.1038/nmeth.2019.

[39] Leda Lacaria, Alessandro Podestà, Manfred Radmacher, and Felix Rico. 2.2 Contact Mechanics. In Malgorzata Lekka, Daniel Navajas, Manfred Radmacher, and Alessandro Podestà, editors, Volume 1 Biomedical Methods, pages 21–64. De Gruyter, February 2023. ISBN 978-3-11-064063-2.

[40] R. E. Mahaffy, C. K. Shih, F. C. MacKintosh, and J. Käs. Scanning probe-based frequency-dependent microrheology of polymer gels and biological cells. Physical Review Letters, 85 (4):880–883, 2000. doi: 10.1103/PhysRevLett.85.880.

[41] Jordi Alcaraz, Lara Buscemi, Mireia Grabulosa, Xavier Trepat, Ben Fabry, Ramon Farré, and Daniel Navajas. Microrheology of human lung epithelial cells measured by atomic force microscopy. Biophysical Journal, 84(3):2071–2079, March 2003. ISSN 0006-3495. doi: 10.1016/S0006-3495(03)75014-0.

[42] Emilios K. Dimitriadis, Ferenc Horkay, Julia Maresca, Bechara Kachar, and Richard S. Chadwick. Determination of elastic moduli of thin layers of soft material using the atomic force microscope. Biophysical Journal, 82(5):2798–2810, May 2002. ISSN 0006-3495. doi: 10.1016/S0006-3495(02)75620-8.

[43] Y. Abidine, V. M. Laurent, R. Michel, A. Duperray, and C. Verdier. Local mechanical properties of bladder cancer cells measured by AFM as a signature of metastatic potential. The European Physical Journal Plus, 130(10):202, October 2015. ISSN 2190-5444. doi: 10.1140/epjp/i2015-15202-6.

[44] Yara Abidine, Andrei Constantinescu, Valérie M. Laurent, Vinoth Sundar Rajan, Richard Michel, Valentin Laplaud, Alain Duperray, and Claude Verdier. Mechanosensitivity of Cancer Cells in Contact with Soft Substrates Using AFM. Biophysical Journal, 114(5):1165–1175, March 2018. ISSN 00063495. doi: 10.1016/j.bpj.2018.01.005.

[45] J. Alcaraz, L. Buscemi, M. Puig-de-Morales, J. Colchero, A. Baró, and D. Navajas. Correction of Microrheological Measurements of Soft Samples with Atomic Force Microscopy for the Hydrodynamic Drag on the Cantilever. Langmuir, 18(3):716–721, February 2002. ISSN 0743-7463, 1520-5827. doi: 10.1021/la0110850.

[46] Kheya Sengupta and Laurent Limozin. Adhesion of Soft Membranes Controlled by Tension and Interfacial Polymers. Physical Review Letters, 104(8):088101, February 2010. doi: 10.1103/PhysRevLett.104.088101.

[47] U. Seifert and R. Lipowsky. Morphology of Vesicles. In Handbook of Biological Physics, volume 1, pages 403–463. Elsevier, 1995. ISBN 978-0-444-81975-8. doi: 10.1016/S1383-8121(06)80025-4.

[48] Reinhard Lipowsky and Udo Seifert. Adhesion of vesicles and membranes. Molecular Crystals and Liquid Crystals, 202(1):17–25, 1991. ISSN 1056-8816. doi: 10.1080/00268949108035656.

[49] E. Evans and Y. C. Fung. Improved measurements of the erythrocyte geometry. Microvascular Research, 4(4):335–347, October 1972. ISSN 0026-2862. doi: 10.1016/0026-2862(72)90069-6.

[50] Gaurav D Bhabhor, Chetna Patel, Nishant Chhillar, Arun Anand, and Kirit N Lad. Geometrical characterization of healthy red blood cells using digital holographic microscopy and parametric shape models for biophysical studies and diagnostic applications. Journal of Physics D: Applied Physics, 57(35):355401, 2024. ISSN 0022-3727, 1361-6463. doi: 10.1088/1361-6463/ad5025.

[51] Karolina Chrabąszcz, Monika Szczepanek-Dulska, Piotr Deptuła, Robert Bucki, and Katarzyna Pogoda. Engineering cell adhesion: Poly-l-lysine-induced micro- and nanoscale surface modifications and their impact on cellular behavior and subcellular spectral signatures. Spectrochimica Acta Part A: Molecular and Biomolecular Spectroscopy, 348:127110, March 2026. ISSN 1386-1425. doi: 10.1016/j.saa.2025.127110.

[52] Shamik Sen, Shyamsundar Subramanian, and Dennis E. Discher. Indentation and Adhesive Probing of a Cell Membrane with AFM: Theoretical Model and Experiments. Biophysical Journal, 89(5):3203–3213, November 2005. ISSN 00063495. doi: 10.1529/biophysj.105.063826.

[53] Sarah Barns, Marie Anne Balanant, Emilie Sauret, Robert Flower, Suvash Saha, and Yuan-Tong Gu. Investigation of red blood cell mechanical properties using AFM indentation and coarse-grained particle method. BioMedical Engineering OnLine, 16(1):140, December 2017. ISSN 1475-925X. doi: 10.1186/s12938-017-0429-5.

[54] Dominic Vella, Amin Ajdari, Ashkan Vaziri, and Arezki Boudaoud. Indentation of Ellipsoidal and Cylindrical Elastic Shells. Physical Review Letters, 109(14):144302, October 2012. ISSN 0031-9007, 1079-7114. doi: 10.1103/PhysRevLett.109.144302.

[55] Edith Schäfer, Marian Vache, Torben-Tobias Kliesch, and Andreas Janshoff. Mechanical response of adherent giant liposomes to indentation with a conical AFM-tip. Soft Matter, 11(22):4487–4495, May 2015. ISSN 1744-6848. doi: 10.1039/C5SM00191A.

[56] Amaury Perez-Tirado, Ulla Unkelbach, and Andreas Janshoff. Tissue tension of planar, free standing cell monolayers measured by central deformation. PNAS Nexus, 4(10):pgaf324, October 2025. ISSN 2752-6542. doi: 10.1093/pnasnexus/pgaf324.

[57] Daan Vorselen, Fred C. MacKintosh, Wouter H. Roos, and Gijs J.L. Wuite. Competition between Bending and Internal Pressure Governs the Mechanics of Fluid Nanovesicles. ACS Nano, 11(3):2628–2636, March 2017. ISSN 1936-0851. doi: 10.1021/acsnano.6b07302.

[58] Ben Fabry, Geoffrey N. Maksym, James P. Butler, Michael Glogauer, Daniel Navajas, Nathan A. Taback, Emil J. Millet, and Jeffrey J. Fredberg. Time scale and other invariants of integrative mechanical behavior in living cells. Physical Review E, 68(4):041914, October 2003. ISSN 1063-651X, 1095-3787. doi: 10.1103/PhysRevE.68.041914.

[59] Bob Fregin, Fabian Czerwinski, Doreen Biedenweg, Salvatore Girardo, Stefan Gross, Konstanze Aurich, and Oliver Otto. High-throughput single-cell rheology in complex samples by dynamic real-time deformability cytometry. Nature Communications, 10(1):415, January 2019. ISSN 2041-1723. doi: 10.1038/s41467-019-08370-3.

[60] Agathe Nidriche, Magalie Faivre, Taha Benyattou, and Claude Verdier. Red blood cells rheology measured using afm. a comparison with other studies. Multidisciplinary Biomechanics Journal, 50th congress of the Société de Biomécanique:101, oct. 2025. ISSN 3076-1158. doi: 10.46298/mbj.16234.

[61] Dmitry A. Fedosov, Huan Lei, Bruce Caswell, Subra Suresh, and George E. Karniadakis. Multiscale Modeling of Red Blood Cell Mechanics and Blood Flow in Malaria. PLoS Computational Biology, 7(12):e1002270, December 2011. ISSN 1553-7358. doi: 10.1371/journal.pcbi.1002270.

[62] Dmitry A. Fedosov, Bruce Caswell, and George Em Karniadakis. A Multiscale Red Blood Cell Model with Accurate Mechanics, Rheology, and Dynamics. Biophysical Journal, 98 (10):2215–2225, May 2010. ISSN 00063495. doi: 10.1016/j.bpj.2010.02.002.

