## Supplemental materials for "Influence of Non-Specific Surface Adhesion on the Shape and Microrheology of Red Blood Cells"

#### 1 Supplementary information to *Materials and Methods*

##### 1.1 PLL coating characterization

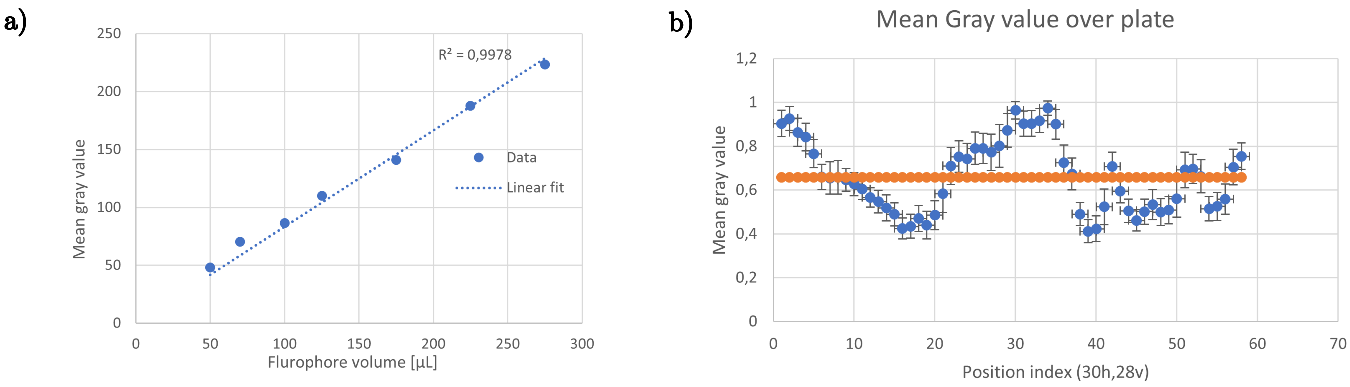


Figure S1: a) Linear trend between the measured intensity in gray levels in $[0, 255]$ and the quantity of sulforhodamine B fluorophore probed by the increase of total volume where $c_{\text{sulforhodamine}} = 5 \times 10^{-4}\,\text{mg/mL}$ (molar stoechiometry for $c_{\text{PLL}} = 10\,\text{mg/mL}$). b) Mean gray value (blue curve) and average mean gray value (orange curve) over the $c_{\text{PLL}} = 10\,\text{mg/mL}$ functionalized glass plate, with 30 frames over the horizontal cross section and 28 along the vertical cross section defined from the center, in steps of $0.5\,\mu\text{m}$.

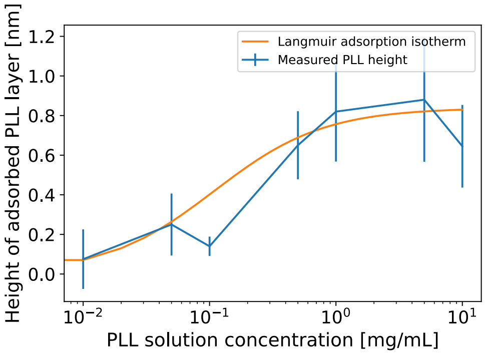


Figure S2: Average height of dry PLL adsorbed on a silicon wafer (10 minutes of incubation time) as a function of the PLL solution concentration $c_{\text{PLL}}$, measured using ellipsometry. Data is featured in blue (lines are a guide for the reader), and the Langmuir adsorption model is displayed in orange

#### 1.2 AFM microrheology measurements

##### 1.2.1 Substrate effect correction

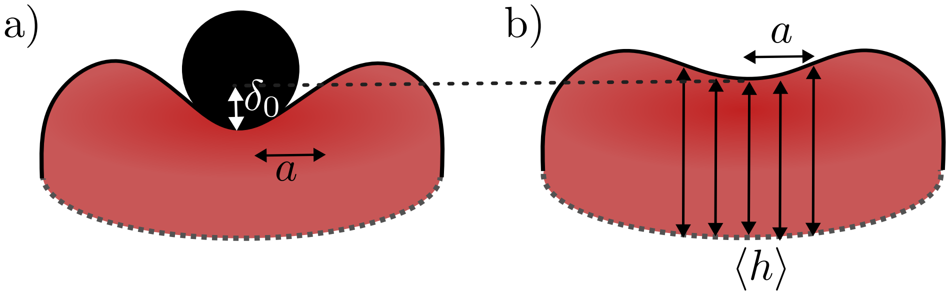


Figure S3: Estimation of the RBC's height from topography curves (AFM). a) Indented RBC, estimation of the radius of contact $a = \sqrt{R^2 - (R - \delta_0)^2}$ for $R > \delta_0$ from Hertz model using indentation $\delta_0$, measured for all RBC. b) Estimation of $\langle h \rangle = \frac{1}{2a} \int_{-a}^{a} h(r)dr$ with r the radial distance from the center of the RBC.

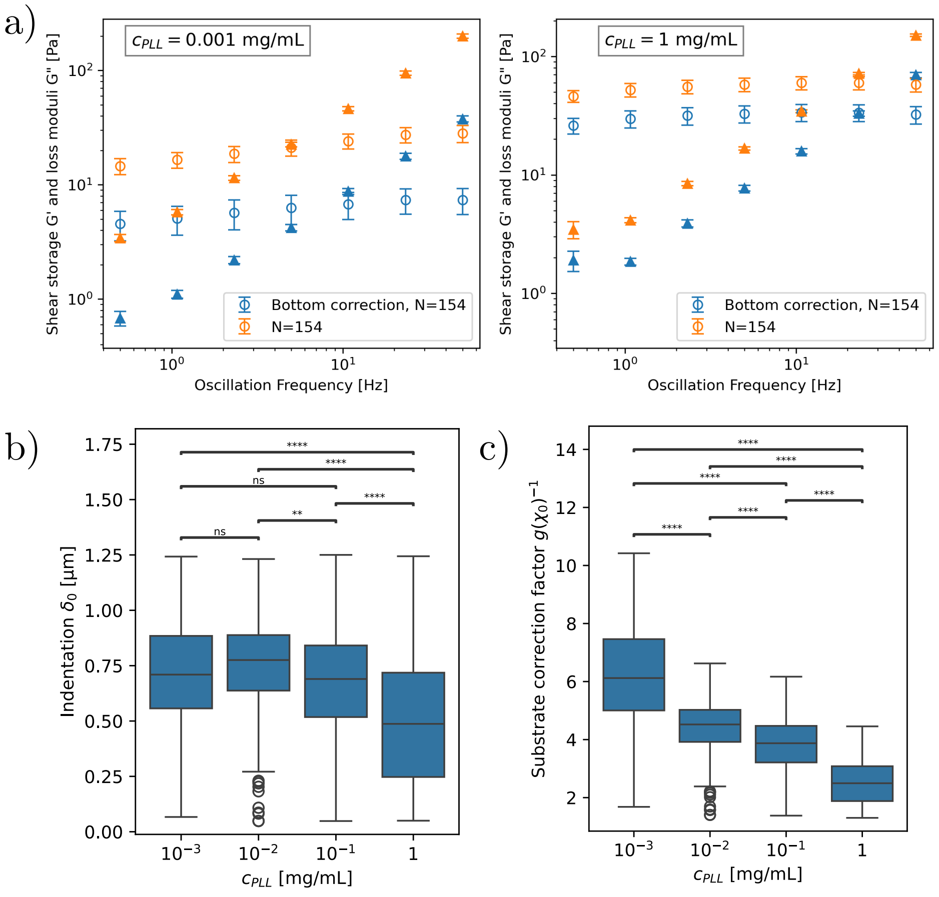


Figure S4: a) Shear $G'(\omega)$ and loss $G"(\omega)$ moduli obtained with (blue curve) and without (orange curve) substrate correction to compensate the effect of the substrate. b) Boxplot of spherical tip indentation $\delta_0$, c) boxplot of the correction factors $g(\chi_0)$ used to correct $E$, $G'(\omega)$ and $G"(\omega)$ for each RBC.

Figure S4 (a) shows that the complex shear modulus $G^*$ (orange curve) is strongly diminished once corrected for the finite thickness of the sample (blue curve) for the two extreme concentrations $c_{\text{PLL}} = 10^{-3}$ mg/mL and $c_{\text{PLL}} = 1$ mg/mL. The correction $g(\chi_0)^{-1} = (1 + 1.511\chi_0 + 2.138\chi_0^2 + 1.583\chi_0^3 + 0.2275\chi_0^4)^{-1}$ where $\chi_0 = \sqrt{R\ \delta_0}/h$ is important due to the large indentation of the sample.

In order to apply the substrate correction, both $\delta_0$ and $h$ are required. Indentation $\delta_0$ is defined as the distance between the absolute position of the contact point defined with Hertz model and the absolute position in the sample where oscillations are performed. $h$ is the height of the undeformed RBC at the center. While $\delta_0$ is precisely estimated for each cell for each measurement, precise estimation of $h$ is unavailable.

Therefore, $\langle h \rangle$ is obtained for each concentration using the QI mode topography images, and is described in Figure S3 (a) and (b). $\langle h \rangle$ is calculated with the central RBC height before indentation and averaged over the median contact radius $a$ of the spherical indenter, such that $\langle h \rangle = \frac{1}{2a} \int_{-a}^{a} h(r)dr$ (column 3 of Table S1). $a = \sqrt{R^2 - (R - \delta_0)^2}$ (column 2 of Table S1) is estimated using the median indentation $\delta_0$ of each concentration (column 1 of Table S1). The distribution of $\delta_0$ for the entire set of data is plotted on Figure S4 (c), and shows that indentation spans a large fraction of the RBC's height at the center. This provides large correction factors plotted on Figure S4 (d), which decrease almost linearly with concentration $c_{\mathrm{PLL}}$.

| $c_{\mathrm{PLL}}$ [mg/mL] | Median indentation $\delta_0$ [$\mu$m] | Median Contact Radius $a$ [$\mu$m] | Average RBC height $\langle h \rangle$ [$\mu$m] |
|---|---|---|---|
| $10^{-3}$ | 0.72 | 1.31 | $1.06 \pm 0.20$ |
| $10^{-2}$ | 0.79 | 1.26 | $1.34 \pm 0.12$ |
| $10^{-1}$ | 0.69 | 1.11 | $1.41 \pm 0.18$ |
| 1 | 0.49 | 1.28 | $1.71 \pm 0.13$ |

Table S1: Average RBC height $\langle h \rangle$ calculated using the median contact radius of the indenter $a$ and the median indentation $\delta_0$ for each concentration $c_{\mathrm{PLL}}$.

##### 1.2.2 Hydrodynamic drag correction

The protocol for hydrodynamic drag correction is explained in the main text. The implementation of the correction follows the methodology of Alcaraz *et al* [1] in a custom-made Python script available in our GitHub repository. It takes the tip and solvent characteristics as inputs (bending rigidity $k$ in N/m, viscosity $\eta$) in order to get the drag transfer function $H_d(f) = \frac{k\frac{F(f)}{z(f)}}{k-\frac{F(f)}{z(f)}}$ relating the imposed sinusoidal displacement $z(f)$ of the cantilever to the measured sinusoidal force $F(f)$ exerted by the solvent.
Since the solvent is purely viscous, the real part $H'(d)$ is supposedly zero and its imaginary part $H_d(f)'' = 2\pi i f b(h)$ is linear with the frequency $f$. We observed the linear trend for $f \leq 200$ Hz on Figure S5 (a), and we verify that the force is dissipative with $H'(d) \approx 0$. The linear regression provides the fit parameter $b(h)$, which is the drag factor at height $h$. $b(h)$ follows $b(h) = 6\pi\eta a_{\text{eff}}^2/(h + h_{\text{eff}})$ which is a drag factor modified from the ideal spherical geometry, introducing an effective area $a_{\text{eff}}$ and an effective height $h_{\text{eff}}$ which eliminates the singularity at $h = 0$. The extrapolation for $h = 0$ provides the drag factor $b(0)$ at the surface of the sample. The fit is shown in Figure S5 (b), and is performed for $h \in [500, 5000]$ nm.
5 consecutive measurements on the same QUEST20 tip with attached 3$\mu$m diameter bead yields a correction factor $b(0) = 3.88 \pm 0.03\,\mu$Ns/m, fit parameters are exposed in Table S2.

| Tip stiffness [N/m] | Tip geometry | Cantilever geometry | b(0) | $a_{\text{eff}}$ [$\mu$m$^2$] | $h_{\text{eff}}$ [$\mu$Ns/m] [$\mu$m] |
|---|---|---|---|---|---|
| 0.02 | 3$\mu$m diameter sphere | Triangular | $3.88 \pm 0.03$ | $6.62 \pm 0.24$ | $25.56 \pm 0.83$ |

Table S2: Fitted drag coefficient at $h = 0$ and fitted parameters $a_{\text{eff}}$ and $h_{\text{eff}}$ of QUEST20 cantilever.

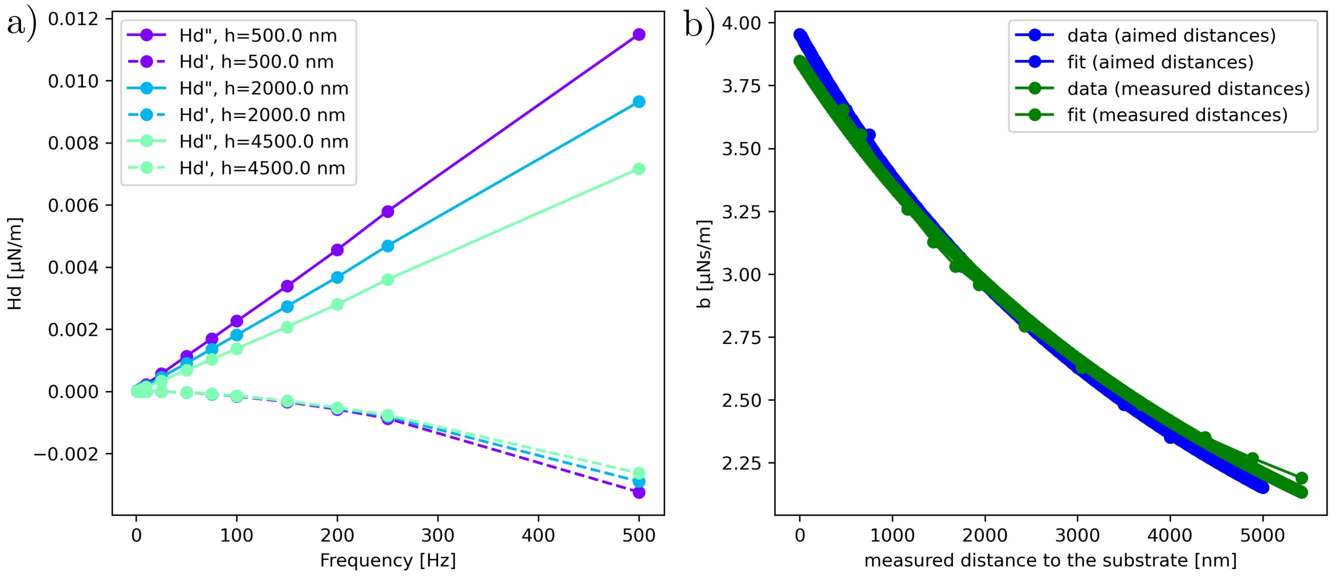


Figure S5: (a) Transfer function $H_d$ divided in its imaginary part $H$"$(d)$ and its real part $H'(d)$ as a function of the sinusoidal frequency $f$ of the imposed tip displacement.(b) Drag coefficient $b(h)$ as a function of substrate-tip height $h$, modeled with the modified sphere model. The blue curve corresponds to the height monitored during measures, and the green curves to the actual corrected relative height of the cantilever.

#### 1.3 Statistical analysis

| Experiment | $N$ | RBCs per $c_{\mathrm{PLL}}$ in mg/mL 0 | $10^{-3}$ | $10^{-2}$ | $10^{-1}$ | $10^{0}$ |
|---|---|---|---|---|---|---|
| RICM, contact area | 1 | 258 | 258 | 515 | 462 | 378 |
| RICM, fluctuations | 1 | 140 | 154 | 101 | 0 | 123 |
| AFM, scans | 3 | 0 | 15 | 15 | 8 | 19 |
| AFM, $E$, $G^*$ ($F = 30\,\mathrm{pN}$) | 11 | 0 | 154 | 143 | 182 | 154 |
| AFM, $E$ ($F = 200\,\mathrm{pN}$) | 2 | 0 | 73 | 56 | 83 | 94 |

Table S3: Number of RBCs measured, depending on PLL concentration. $N$ corresponds to the number of donors.

### 2 Supplementary information to *Non-specific cell-substrate adhesion triggers a transition in RBC geometry*

#### 2.1 Confocal microscopy

Confocal microscopy experiments corroborate the observation of dome-shaped RBC at $c_{\mathrm{PLL}} = 1\,\mathrm{mg/mL}$ with AFM and RICM. The labeling of the glycocalyx is performed using Alexa red fluorophore 488 (ThermoFisher Scientific, Waltham Massachusetts, U.S.), using a laser wavelength $\lambda = 488\,\mathrm{nm}$ and a $63\times$ oil objective. Figure S6 displays a $x$-$y$ plot of a labeled RBC at the contact of the substrate and 2 perpendicular $z$-projections of the RBC with $\Delta z \approx 0.5\,\mu\mathrm{m}$.

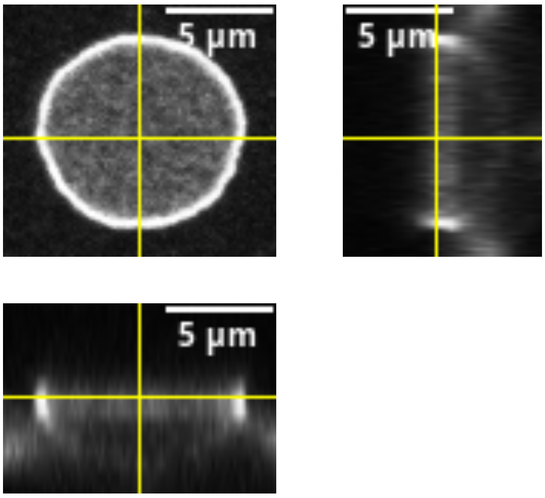


Figure S6: Confocal microscopy image of a single RBC performed with $c_{\mathrm{PLL}} \approx 1\,\mathrm{mg/mL}$, the orthogonal slices show a probable spherical cap shape of the RBC.

#### 2.2 RICM data: evolution of the shape of the RBC

It appears clearly that with increasing concentration, the shape of cells evolves from a more rounded, even echinocyte-like shape as reported by Hategan et al [2], to a tensed elliptical shape.

We extract two uncorrelated shape descriptors (corr$<$ 0.8) provided in ImageJ software: Solidity ($S = A^{*}/\text{convex}$ $A*$) providing a description of the convexity of the adhesion disk, and Round ($R = 4\pi A/2\pi a$ with $a$ the major ellipse axis) providing a description of the anisotropy of the adhesion disk. As observed on Figure S7 (typical images are featured below the graphs), $S$ shows that the convexity of the RBC clearly increases with $c_{\text{PLL}}$ ($S$), while $R$ shows that it gets also more oriented due to the presence of poly-L-lysine.

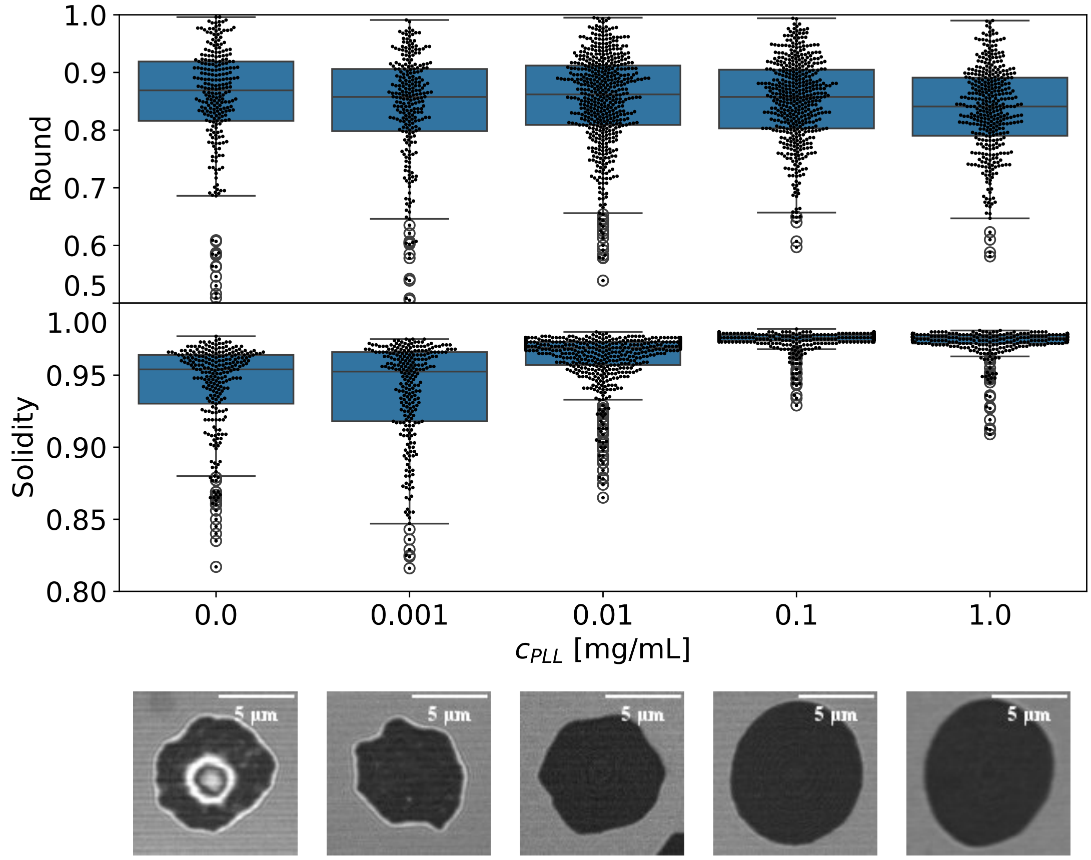


Figure S7: Bottom surface description of the RBC: Round $R$ and Solidity $S$ shape descriptors as a function of $c_{\text{PLL}}$, $N = [258, 258, 515, 462, 378]$. The bottom images ($10.81\,\mu\text{m} \times 10.81\,\mu\text{m}$) were obtained with RICM and illustrate the distribution of RBC adhesion profiles for each $c_{\text{PLL}}$ concentration.

#### 2.3 AFM QI mode

| $c_{\mathrm{PLL}}$ [mg/mL] | $R_0$ (Spreading radius $\mu$m) | | | $C_0$ (thickness $\mu$m) | | | $C_2$ $\mu$m | | | $C_4$ $\mu$m | | |
|---|---|---|---|---|---|---|---|---|---|---|---|---|
| | Conc. | Flat | Dome | Conc. | Flat | Dome | Conc. | Flat | Dome | Conc. | Flat | Dome |
| $10^{-3}$ | 4.7 | 4.7 | 5.1 | 0.7 | 1.4 | 1.2 | 2.1 | 2.1 | 2.1 | −2.8 | −3.5 | −3.9 |
| $10^{-2}$ | 4.9 | / | 4.9 | 1.1 | / | 1.6 | 2.5 | / | 1.1 | −2.7 | / | −3.4 |
| $10^{-1}$ | / | 4.9 | 5.1 | / | 1.1 | 1.8 | / | 1.5 | −0.4 | / | −1.5 | −1.3 |
| 1 | / | 4.9 | 5.2 | / | 1.3 | 1.8 | / | 1.1 | −0.5 | / | −1.9 | −1.3 |
| Fung | 3.91 | | | 0.81 | | | 7.83 | | | −4.39 | | |

Table S4: Table of fitting parameters for the RBC's shape obtained with AFM and compared to native RBC parameters reported by Fung *et al.* [3], for different $c_{\mathrm{PLL}}$ concentrations.

.

| Cluster # | $R_0$ [$\mu$m] (radius $R^*$) | $C_0$ [$\mu$m]) (thickness) | $C_2$ [$\mu$m] | $C_4$ [$\mu$m] |
|---|---|---|---|---|
| 1: concave | 4.9 | 0.8 | 3.4 | −4.2 |
| 2: flat | 4.9 | 1.3 | 1.1 | −2.1 |
| 3: dome | 5.1 | 2.1 | −0.9 | −0.1 |
| Fung | 3.91 | 0.81 | 7.83 | −4.39 |

Table S5: Table of mean values obtained in each cluster for the parameters of the Fung's model for RBCs geometry, eq. 8.

The distribution of the substrate effect corrected Young's moduli $E_c$ obtained using the quantitative Imaging (QI) mode using the sharp cylindrical tip of radius 70 nm of the PFQNM-LC-V2 probe with force setpoint $F \approx 40$ pN is shown on Figure S8, $N = [15, 15, 8, 19]$. Although values are insufficiently rigorous due to the specific shape of this tip optimized for imaging (paraboloid, conical and cylindrical depending on the indentation depth), they corroborate that only the dome shaped RBCs from cluster #3 are widely dispersed towards high $E$ values, while concave and flat RBCs from clusters #1 and #2 are similar in mechanical properties. The dependence of $E$ on the tip shape is typical of AFM measurement, whether a local probe (PFQNM-LC-V2) or a global probe (QUEST T20) is being used [4].

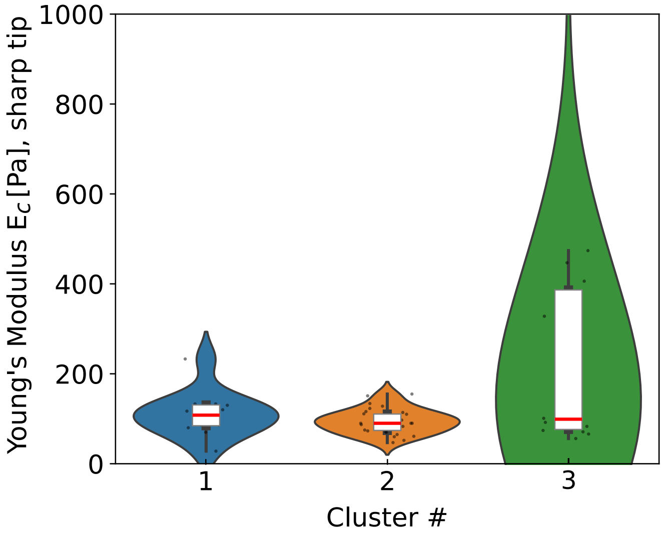


Figure S8: Young Moduli $E$ obtained using Bruker QI mode with PFQNM-LC-V2 probe and corrected for the substrate stiffness for a cylindrical geometry. Data is sorted into the clusters found in the topography section.

As explained in the Results section, imaging soft live cells in contact mode is complicated by the ill-determination of the contact point, appearing through an under-estimation of the RBC's height. The error in contact-point determination is estimated assuming that the volume of the RBC is constant from the lowest to the highest PLL concentrations, and that the error in contact point is uniform across the RBC. Averaging over references [5–7], the typical volume of a healthy adult RBC is found around $V_0 \approx 84\ \mu\text{m}^3$. Introducing a height offset $h_{\text{offset}}$ verifying $V(x, y, z + h_{\text{offset}}) = V_0$, we observe that $h_{\text{offset}}$ decreases with $c_{\text{PLL}}$ from 400 to 30 nm, since the contact point and therefore the height are better determined for tensed RBC. Furthermore, the tip is sharp (70 nm in radius), which is necessary for topography mapping but implies a deeper indentation into the cell than for larger radii.

| Concentration $c_{\text{PLL}}$ [mg/mL] | 0.001 | 0.01 | 0.1 | 1 |
|---|---|---|---|---|
| RBC contact area $A^*$ [$\mu$m$^2$] | 48 | 53 | 61 | 62 |
| RBC volume $V$ [$\mu$m$^3$] | 56 | 69 | 71 | 76 |
| $h_{\text{offset}}$ [$\mu$m], $V(x, y, z + h_{\text{offset}}) = V_0$ | 0.39 | 0.14 | 0.05 | 0.03 |

Table S6: RBC cross-section area $A^*$ and volume $V$ obtained with AFM. $h_{\text{offset}}$ is the error in contact-point determination.

#### 3 Supplementary information to *A clear limitation of the Hertz model*

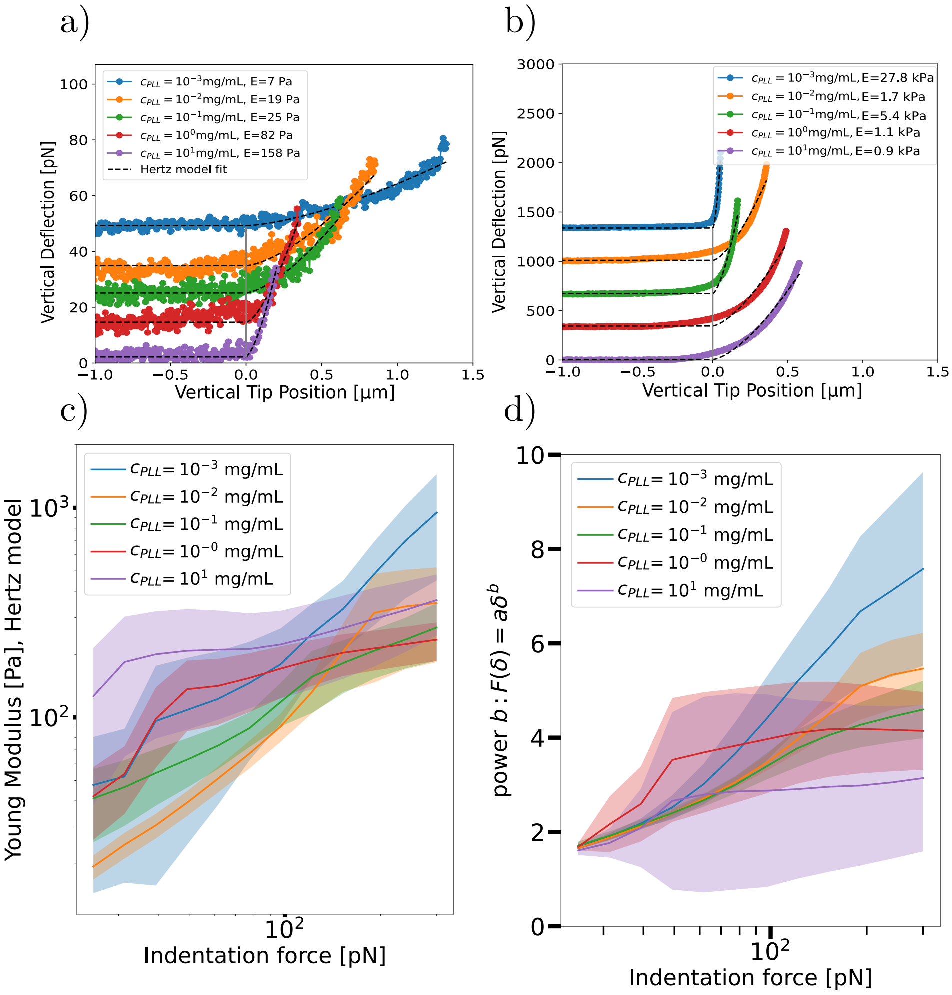


Figure S9: Study of the limitation of Hertz model for the study of RBCs mechanical properties. A force offset is applied for better visualisation of force curves. a) and b), force curves of 5 RBCs at different $c_{\mathrm{PLL}}$ incubation concentrations in $[10^{-3}, 10^{-2}, 10^{-1}, 10^{0}, 10^{1}]$ mg/mL obtained for setpoints $F = 30$ pN (a) and $F = 1000$ pN (b), fitted using the Hertz model (black dotted lines). c) Young's Modulus $E$ obtained with Hertz model for varying force setpoints $F$ referred to as "indentation force" , plotted as the mean $\pm 95\%$ confidence interval. d) Best fitted power-law exponent to the force-indentation curves for varying force setpoint $F$. $N = [73, 56, 83, 94, 12]$

### 4 Supplementary information to *The rheology of the RBC is impacted by the adhesion strength to the substrate*

#### 4.1 Rheology models comparison

Figure S10 compares the shear storage and loss moduli, respectively $G'$ (exp data: open dots) and $G''$ (exp data: closed triangles), obtained using the structural damping model (a) and the Kelvin Voigt model (b). It also compares the trends obtained for the loss tangent corresponding to $G''/G'$. The model is represented using continuous lines, matching the colour-code of experimental points. While both models capture similarly the experimental points at high PLL concentration, the structural damping model fits better the data at low concentration. More specifically, in the low frequency region which is also known to capture the ATP activity of the RBC [8].

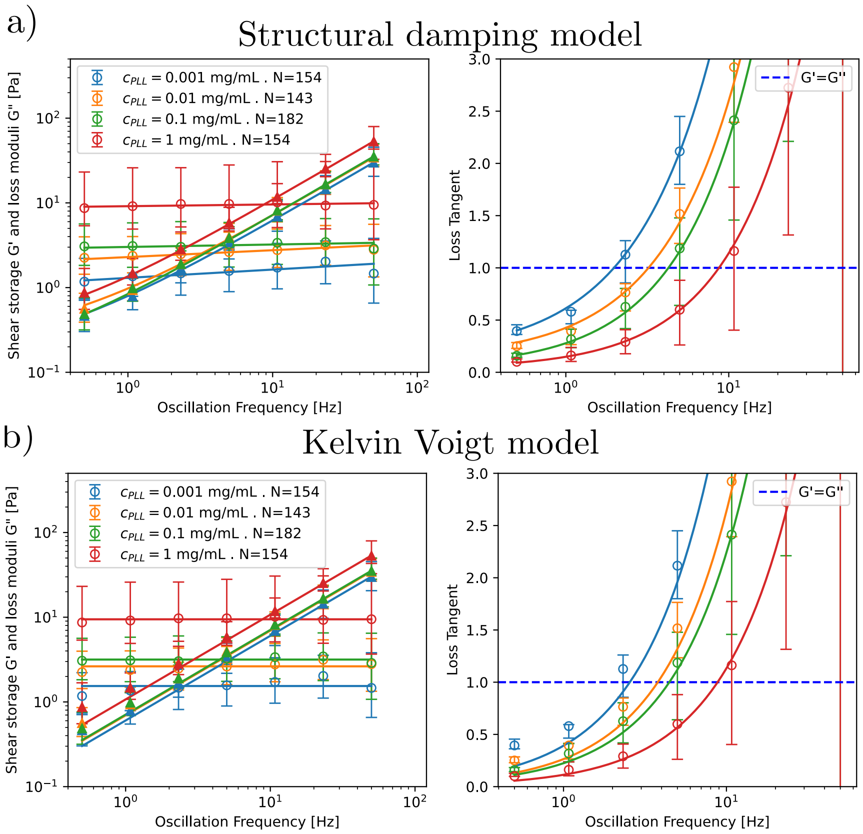


Figure S10: Comparison of the structural damping model to the Kelvin-Voigt model, which is a specific case of the first model where $\alpha = 0$ (see equation 9, main text).

| Authors | Method | $G_0$ [Pa] (storage) | Power-law exponent $n$ (loss) |
|---|---|---|---|
| Bergamaschi *et al.* [9] | AFMR | $72 \pm 4$ | $0.47 \pm 0.05$ |
| Puig de Morales *et al.* [10] | MTC | | 0.64 |
| Gomez *et al.* [11] | Optical Tweezers | $49 \pm 5$ | $0.65 \pm 0.06$ |
| Amin *et al.* [12] | Dynamic light scattering | $[0.01, 0.1]$ | 0.69 |
| Current study | AFM | $8.9 \pm 1.45$ | $0.93 \pm 0.01$ |

Table S7: Comparison of $G^*$ data with known results, using a simple power-law model $G''(\omega) \propto \omega^n$. Our data is averaged on all PLL concentrations, mean±s.e.m.

#### 4.2 Distribution of parameters for varying blood donors

8 blood donors with comparable measurement statistics are compared on Figure S11. The fit parameters obtained with the power law model adjusted on the microrheology data are plotted for the minimal concentration ($c_{\mathrm{PLL}} = 10^{-3}$ mg/mL, 1% plate coverage) and the maximal concentration ($c_{\mathrm{PLL}} = 1$ mg/mL, 90% plate coverage) of Poly-L-Lysine. The exact cause of inter-donor variability cannot be unravelled: PLL uniformity, donor variability, storage conditions all contribute to this variability. However, the difference in parameters from low to high concentration is systematic. $G_0$ increases and the distribution tends to widen. As for $\alpha$ exponent, the decrease to $\alpha = 0$ shows that the RBC tends to follow a simple Kelvin-Voigt model for strong adhesion. $\mu$, associated with the membrane and cytosolic viscosity, also increases.

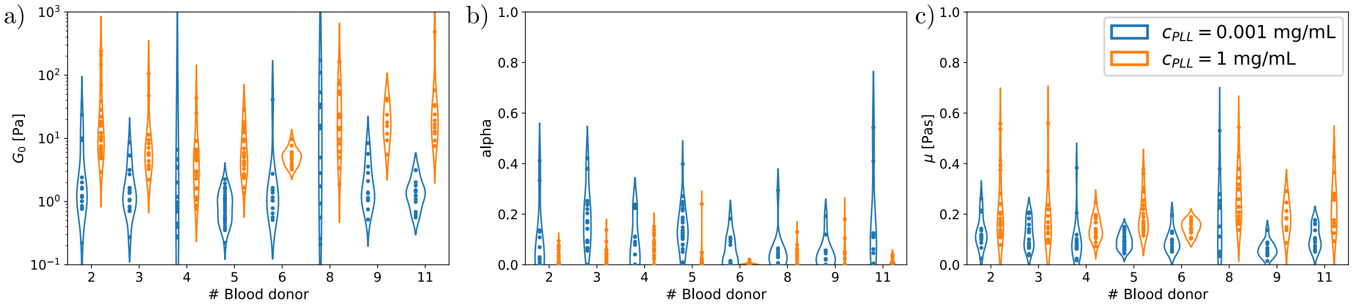


Figure S11: Distribution of fit parameters obtained for all donors and compared for the lowest ($c_{\mathrm{PLL}} = 10^{-3}$ and highest ($c_{\mathrm{PLL}} = 1$mg/mL) concentration of PLL. a) The shear modulus at 0 frequency $G_0$, b) the exponent $\alpha$ of the power-law model, c) Newtonian viscosity $\mu$.

#### 5 Comparison with literature

Table S8: Literature review of RBC Young's modulus $E$ measured using AFM or other scanning methods, comparison of protocols featuring the force setpoint used (and fitting force if different), tip characteristics, and PLL coating protocol.

| Author | Young's Modulus $E$ | $F_{\mathbf{setpoint}}$ | Tip | Plate preparation | Model |
|---|---|---|---|---|---|
| Li *et al.* [13] | $145 \pm 60$ Pa | $F_{\text{fit}} = 200$ pN | 15 $\mu$m diam. spherical indenter | PLL coating | Hertz model |
| Baier *et al.* [14] | 179 Pa (median) | $F_{\text{max}} = 1$ nN, $F_{\text{fit}} \approx 100$ pN | Tetrahedral indenter, $r < 10$ nm, $k = 0.085$ N/m | 0.05 mg/mL PLL (50 $\mu$L), 10 min incubation | Hertz model |
| Ciasca *et al.* [15] | 1.87 kPa (edges: 60 Pa, center: 9 kPa) | $F_{\text{max}} = 500$ pN | 40° conical indenter $r \approx 10$ nm, $k = 0.05$ N/m | PLL coating | Hertz model |
| Wang *et al* [16] | $2.6 \pm 1.2$ kPa | $F_{\text{max}} = 1$ nN, $F_{\text{fit}} = 80$ pN | $r < 10$ nm, $k = 0.06$ N/m | PLL coating | Hertz model |
| Lekka *et al.* [17] | $4.9 \pm 0.5$ kPa | $F_{\text{max}} \approx 6$ nN | $r = 50$ nm, $k = 0.01$ N/m | 0.1 mg/mL PLL coating, 5 min incubation | Sneddon Model |
| Barns *et al.* [18] | $7.57 \pm 3.25$ kPa | $F_{\text{fit}} \in [0.5 - 2.5\,\text{nN}]$ | 5 $\mu$m diam. spherical indenter | 0.1 mg/mL Poly-D-Lysine coating, 10 min incubation + 1% glutaraldehyde | Hertz, substrate correction |
| Sergunova *et al.* [19] | $10 \pm 4$ kPa | $F_{\text{max}} = 60$ nN | Paraboloid indenter, $r = 10$ nm, $k = 1$ N/m | PLL coating | Sneddon model |
| Dulinska *et al.* [20] | $26 \pm 7$ kPa | $F_{\text{max}} \approx 2$ nN | $r = 45$ nm, 0.03 N/m | 1 mg/mL PLL coating + 0.5% glutaraldehyde | Sneddon model |
| Girasole *et al.* [21] | $98 \pm 7$ kPa | $F\text{max} = 2.5$ nN | Pyramidal indenter, $r = 10$ nm, $k \in [0.03 - 0.05]$ N/m | PLL coating | Bilodeau model |
| Tognoni *et al.* [22] | $E \in [0.2 - 1.5]$ kPa | $F_{\text{max}} \in [20 - 500]$ pN | Scanning Ion Conductance Microscopy | 0.1 mg/mL PLL coating, 30 min incubation | / |

#### References


[1] Jordi Alcaraz, Lara Buscemi, Mireia Grabulosa, Xavier Trepat, Ben Fabry, Ramon Farré, and Daniel Navajas. Microrheology of human lung epithelial cells measured by atomic force microscopy. *Biophysical Journal*, 84(3):2071–2079, March 2003.

[2] Alina Hategan, Kheya Sengupta, Samuel Kahn, Erich Sackmann, and Dennis E. Discher. Topographical Pattern Dynamics in Passive Adhesion of Cell Membranes. *Biophysical Journal*, 87(5):3547–3560, November 2004.

[3] E. Evans and Y. C. Fung. Improved measurements of the erythrocyte geometry. *Microvascular Research*, 4(4):335–347, October 1972.

[4] Shruti G. Kulkarni, Sandra Pérez-Domínguez, and Manfred Radmacher. Influence of cantilever tip geometry and contact model on AFM elasticity measurement of cells. *Journal of Molecular Recognition*, 36(7):e3018, 2023.

[5] Gaurav D Bhabhor, Chetna Patel, Nishant Chhillar, Arun Anand, and Kirit N Lad. Geometrical characterization of healthy red blood cells using digital holographic microscopy and parametric shape models for biophysical studies and diagnostic applications.

[6] Donald N. Houchin, John I. Munn, and Benjamin L. Parnell. A Method for the Measurement of Red Cell Dimensions and Calculation of Mean Corpuscular Volume and Surface Area. *Blood*, 13(12):1185–1191, December 1958.

[7] Otwin Linderkamp, Paul Y. K. Wu, and Herbert J. Meiselman. Geometry of Neonatal and Adult Red Blood Cells. *Pediatric Research*, 17(4):250–253, April 1983.

[8] H. Turlier, D. A. Fedosov, B. Audoly, T. Auth, N. S. Gov, C. Sykes, J.-F. Joanny, G. Gompper, and T. Betz. Equilibrium physics breakdown reveals the active nature of red blood cell flickering. *Nature Physics*, 12(5):513–519, May 2016.

[9] Giulia Bergamaschi, Kees-Karel H. Taris, Andreas S. Biebricher, Xamanie M. R. Seymonson, Hannes Witt, Erwin J. G. Peterman, and Gijs J. L. Wuite. Viscoelasticity of diverse biological samples quantified by acoustic force microrheology (afmr). *Communications Biology*, 7(1):1–14, 2024.

[10] Marina Puig-De-Morales, Mireia Grabulosa, Jordi Alcaraz, Joaquim Mullol, Geoffrey N. Maksym, Jeffrey J. Fredberg, and Daniel Navajas. Measurement of cell microrheology by magnetic twisting cytometry with frequency domain demodulation. *Journal of Applied Physiology*, 91(3):1152–1159, September 2001.

[11] Fran Gómez, Leandro S. Silva, Glauber Ribeiro De Sousa Araújo, Susana Frases, Ana Acacia S. Pinheiro, Ubirajara Agero, Bruno Pontes, and Nathan Bessa Viana. Effect of cell geometry in the evaluation of erythrocyte viscoelastic properties. *Physical Review E*, 101(6):062403, 2020.

[12] M. Shahrooz Amin, YougKeun Park, Niyom Lue, Ramachandra R. Dasari, Kamran Badizadegan, Michael S. Feld, and Gabriel Popescu. Microrheology of red blood cell membranes using dynamic scattering microscopy. *Optics Express*, 15(25):17001, 2007.

[13] Mi Li, LianQing Liu, Ning Xi, YueChao Wang, ZaiLi Dong, XiuBin Xiao, and WeiJing Zhang. Atomic force microscopy imaging and mechanical properties measurement of red blood cells and aggressive cancer cells. *Science China Life Sciences*, 55(11):968–973, November 2012.

[14] Dina Baier, Torsten Müller, Thomas Mohr, and Ursula Windberger. Red Blood Cell Stiffness and Adhesion Are Species-Specific Properties Strongly Affected by Temperature and Medium Changes in Single Cell Force Spectroscopy. *Molecules*, 26(9):2771, January 2021.

[15] G. Ciasca, M. Papi, S. Di Claudio, M. Chiarpotto, V. Palmieri, G. Maulucci, G. Nocca, C. Rossi, and M. De Spirito. Mapping viscoelastic properties of healthy and pathological red blood cells at the nanoscale level. *Nanoscale*, 7(40):17030–17037, October 2015.

[16] Kun Wang, Zhiqiang Li, Ogechukwu Egini, Raj Wadgaonkar, Xian-Cheng Jiang, and Yong Chen. Atomic force microscopy reveals involvement of the cell envelope in biomechanical properties of sickle erythrocytes. *BMC Biology*, 21(1):31, February 2023.

[17] Malgorzata Lekka, Maria Fornal, Grażyna Pyka-Fościak, Kateryna Lebed, Barbara Wizner, Tomasz Grodzicki, and Jan Styczeń. Erythrocyte stiffness probed using atomic force microscope. *Biorheology: The Official Journal of the International Society of Biorheology*, 42(4):307–317, July 2005.

[18] Sarah Barns, Marie Anne Balanant, Emilie Sauret, Robert Flower, Suvash Saha, and YuanTong Gu. Investigation of red blood cell mechanical properties using AFM indentation and coarse-grained particle method. *BioMedical Engineering OnLine*, 16(1):140, December 2017.

[19] Viktoria Sergunova, Stanislav Leesment, Aleksandr Kozlov, Vladimir Inozemtsev, Polina Platitsina, Snezhanna Lyapunova, Alexander Onufrievich, Vyacheslav Polyakov, and Ekaterina Sherstyukova. Investigation of Red Blood Cells by Atomic Force Microscopy. *Sensors (Basel, Switzerland)*, 22(5):2055, March 2022.

[20] Ida Dulińska, Marta Targosz, Wojciech Strojny, Małgorzata Lekka, Paweł Czuba, Walentyna Balwierz, and Marek Szymoński. Stiffness of normal and pathological erythrocytes studied by means of atomic force microscopy. *Journal of Biochemical and Biophysical Methods*, 66(1-3):1–11, March 2006.

[21] Marco Girasole, Simone Dinarelli, and Giovanna Boumis. Structure and function in native and pathological erythrocytes: A quantitative view from the nanoscale. *Micron*, 43(12):1273–1286, December 2012.

[22] Elisabetta Tognoni, Paolo Orsini, and Mario Pellegrino. Nonlinear indentation of single human erythrocytes under application of a localized mechanical force. *Micron*, 127:102760, December 2019.
